# Transferrin receptor 1-mediated iron uptake regulates bone mass in mice via osteoclast mitochondria and cytoskeleton

**DOI:** 10.1101/2021.09.12.459964

**Authors:** Bhaba K Das, Lei Wang, Toshifumi Fujiwara, Jian Zhou, Nukhet Aykin-Burns, Kimberly J Krager, Renny Lan, Samuel G Mackintosh, Ricky Edmondson, Michael L Jennings, Xiaofang Wang, Jian Q Feng, Tomasa Barrientos, Jyoti Gogoi, Aarthi Kannan, Ling Gao, Weirong Xing, Subburaman Mohan, Haibo Zhao

## Abstract

Increased intracellular iron spurs mitochondrial biogenesis and respiration to satisfy high-energy demand during osteoclast differentiation and bone-resorbing activities. Transferrin receptor 1 (TFR1) mediates cellular iron uptake through endocytosis of iron-loaded transferrin and its expression increases during osteoclast differentiation. Nonetheless, the precise functions of TFR1 and TFR1-mediated iron uptake in osteoclast biology and skeletal homeostasis remain incompletely understood. To investigate the role of TFR1 in osteoclast lineage cells, we conditionally deleted *Tfr1* gene in myeloid precursors or mature osteoclasts by crossing *Tfr1*- floxed mice with *LysM-Cre* and *Ctsk*-Cre mice, respectively. Skeletal phenotyping by µCT and histology unveiled that loss of Tfr1 in osteoclast progenitor cells resulted in a three-fold increase in trabecular bone mass in the long bones of 10-week old female but not male mice. Although high trabecular bone volume in long bones was seen in both male and female mice with deletion of *Tfr1* in mature osteoclasts, this phenotype was more pronounced in female knockout mice. Mechanistically, disruption of Tfr1 expression attenuated mitochondrial metabolism and cytoskeletal organization in mature osteoclasts, leading to decreased bone resorption with no impact on osteoclastogenesis. These results indicate that Tfr1-mediated iron uptake is specifically required for osteoclast function and is indispensable for bone remodeling.

## Introduction

Bone mass and structure in adults are maintained by constant bone remodeling with balanced bone formation by osteoblasts and bone resorption by osteoclasts (Zaidi, 2007; Raggatt and Partridge, 2010). Under pathological conditions, however, excessive bone resorption caused by either increased number or exaggerated activities of osteoclasts leads to bone loss which is a hallmark of many metabolic bone diseases such as osteoporosis, rheumatoid arthritis, Paget’s disease of bone, periodontal disease, and tumor bone metastasis (Mbalaviele et al, 2017; Novack and Teitelbaum, 2008). Osteoclasts are multinucleated cells formed by fusion of mononuclear precursors that are differentiated from the monocyte/macrophage lineage of myeloid hematopoietic cells (Walker, 1975; Xiao et al, 2017). The signaling cascades triggered by M-CSF (macrophage colony-stimulating factor) and RANKL (receptor activator of NF-κB ligand) together with intracellular calcium oscillation stimulated by immunoglobulin-like receptors eventually induce and activate NFATc1 (nuclear factor of activated T cells 1), a master transcription factor of osteoclast differentiation (Boyle et al, 2003; Nakashima et al, 2012). Upon attachment to bone, osteoclasts organize their actin cytoskeleton to form actin-rings that seal the resorptive microenvironment (Väänänen et al, 2000; Teitelbaum, 2011). The dissolution of bone minerals and digestion of organic bone matrix mainly type I collagen are executed by hydrochloric acid and acidic hydrolase cathepsin K, respectively, via lysosome secretion (Zhao, 2012; Fujiwara et al, 2016). Both osteoclast formation and bone-resorbing activity of mature osteoclasts demand high energy. However, the pathways and molecular mechanisms regulating osteoclast energy metabolism in osteoclasts remain largely unknown.

Both glycolysis and mitochondrial oxidative phosphorylation (OXPHOS) increase during osteoclast differentiation (Kubatzky et al, 2018; Arnett and Orriss, 2018; Kim et al, 2007; Indo et al, 2013; Li et al, 2020). Osteoclasts contain numerous mitochondria (Chuan, 1931; Holtrop and King, 1977). The mitochondrial respiratory complex I and the key mitochondria transcriptional regulator PGC1β (peroxisome proliferator-activated receptor γ coactivator 1β) are crucial for osteoclast differentiation (Jin et al, 2014; Ishii et al, 2009). Mitochondrial ROS (reactive oxygen species) stimulates osteoclast differentiation (Srinivasan et al, 2010; Kim et al, 2013; Kim H et al, 2017). On the contrary, it has been reported that decreased mitochondrial biogenesis and activity by PGC-1β-deletion in osteoclast lineage cells disrupt osteoclast cytoskeletal organization and function but not osteoclastogenesis (Zhang et al, 2018). Loss of G-protein Gα13 in osteoclast progenitors promotes mitochondrial respiration and cytoskeleton organization but has little effects on osteoclastogenesis in cultures and in mice (Nakano et al, 2019). Heretofore, the role of energy metabolism in osteoclast differentiation and function remain unclear and controversial.

Iron is a nutritional element that plays a fundamental role in mitochondrial metabolism and the biosynthesis of heme and Fe-S clusters which are critical components of the mitochondrial respiratory complexes (supplemental Figure 1A) (Xu et al, 2013). Mammalian cells acquire iron through the uptake of transferrin (Tf), heme, ferritins, and non-transferrin bound iron (NTBI) (supplemental Figure 1B) (Pantopoulos et al, 2012). In Tf-dependent pathway, ferric iron (Fe^3+^)- loaded Tf (holo-Tf) binds to Tf receptors (TFRs) on cell surface and the complex is then internalized via endocytosis. Fe^3+^ is released from Tf in endosomes and is reduced to ferrous iron (Fe^2+^) by the STEAP (six-transmembrane epithelial antigen of the prostate) family of metaloreductases before being transported to the cytoplasm via DMT1 (divalent metal transporter 1) (Gammella et al, 2017). Cellular iron that is not utilized is either stored in ferritins or is exported via ferroportin (FPN) (Donovan et al, 2006; Hentze et al, 2010).

**Figure 1.**
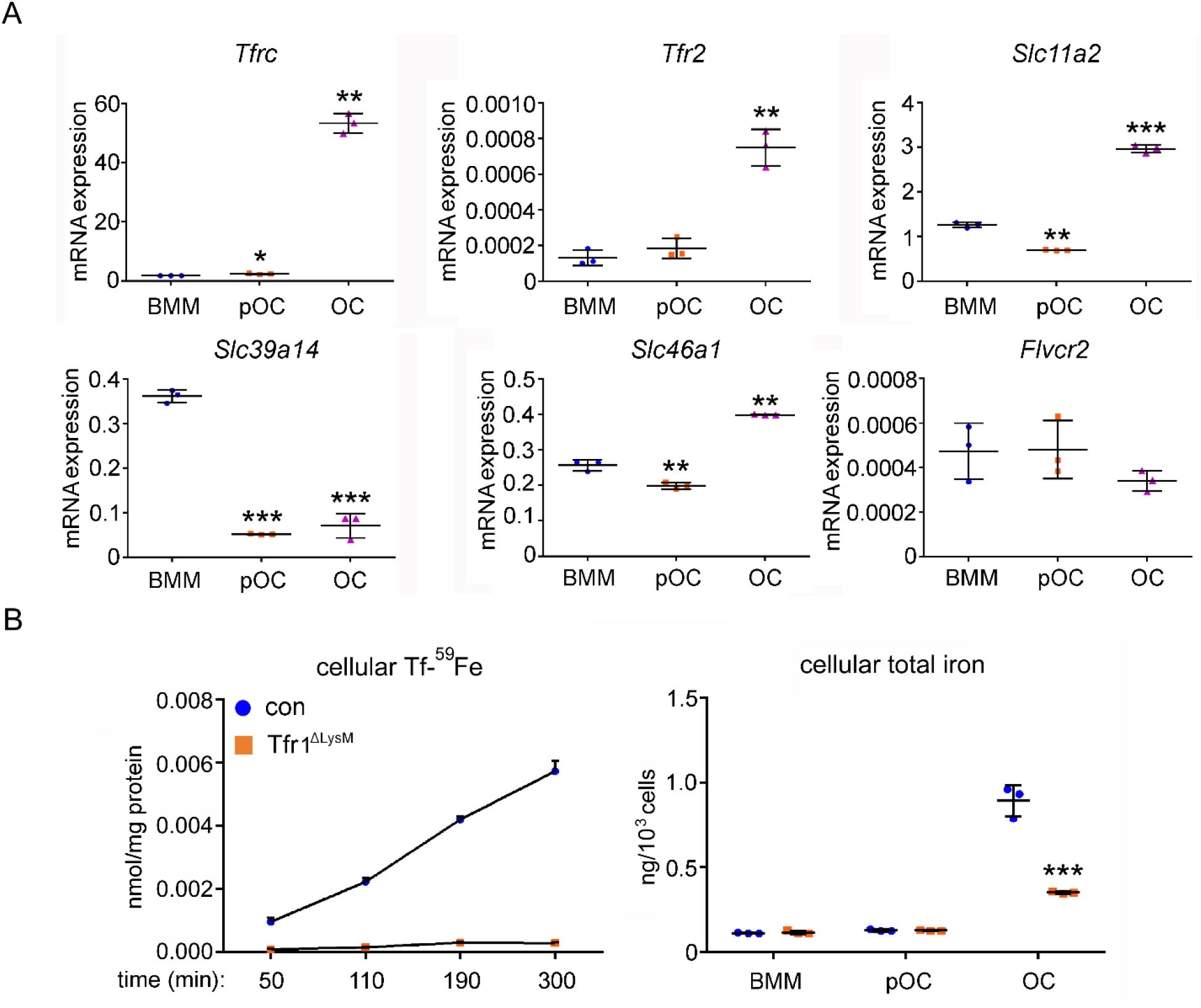
Tfr1 is a major transferrin transporter and regulates cellular iron homeostasis in osteoclasts. (**A**) Detection of mRNA expression of genes involved in iron uptake and export pathways in bone marrow monocytes (BMM), mono-nuclear pre-osteoclasts (pOC), and mature osteoclasts (OC) by real-time quantitative PCR. *Tfrc* encodes Tfr1; *Slc11a1* encodes DMT1 (divalent metal ion transporter 1); *Slc39a14* encodes Zip14; *Slc46a1* encodes Heme transporter HCP1. (**B**) Measurement of ^59^Fe-labeled transferrin (Tf-^59^Fe) uptake in control (con) and Tfr1 myeloid conditional knockout (Tfr1^ΔLysM^) osteoclasts by a gamma counter. (**C**) Measurement of intracellular total iron by a colorimetric iron assay kit in control and Tfr1-deficient osteoclast lineage cells. * p < 0.05, ** p < 0.01, *** p < 0.001 vs BMM (**A**) and vs control OC (**C**) by one-way ANOVA and *Student’s t-test*.

Both systemic and cellular iron homeostasis play a pivotal role in bone remodeling (Balogh et al, 2018). Osteoporosis and pathological bone fractures are commonly associated with the hereditary iron-overload disease hemochromatosis and the acquired iron-overload conditions in treatment of thalassemia and sickle cell disease (Jandl et al, 2020; Almeida A and Roberts, 2005; Dede et al, 2016). Moreover, increased bone resorption and/or decreased bone formation have been observed in genetic mouse models of iron overload diseases and in mice fed with excessive iron (Tsay et al, 2010; Guggenbuhl et al, 2011; Xiao et al, 2015). We have previously reported that excess cellular iron caused by Fpn-deletion in murine myeloid cells stimulates mitochondria and promotes osteoclast number *in vitro* and *in vivo* (Wang et al, 2018). In contrast to iron overload, much less is known about the influence of iron-deficiency on bone cells and bone homeostasis. Severe iron deficiency impairs both bone resorption and bone formation and causes osteopenia in rats (Katsumata et al, 2009; Díaz-Castro et al, 2012), whereas iron chelation inhibits osteoclastogenesis and bone resorption *in vitro* and *in vivo* and attenuates estrogen-deficiency induced bone loss in mice (Ishii et al, 2009; Guo et al, 2015).

There are two TFRs in mammalian cells (Kawabata, 2019). TFR1, encoded by *TFRC* gene, is ubiquitously expressed with high affinity to holo-Tf. Germline deletion of *Tfrc* gene in mice leads to early embryonic lethality due to severe defects in erythroid and neuronal development (Levy et al, 1999). A missense mutation in *TFRC* causes combined immunodeficiency in human (Jabara et al, 2016). The specific deletion of *Tfrc* gene in neurons, skeletal muscles, cardiomyocytes, hematopoietic stem cells, and adipocytes in mice disrupts cellular iron homeostasis and cause severe metabolic defects in these tissues (Matak et al, 2016; Barrientos et al, 2015; Xu et al, 2015; Wang et al, 2020; Li et al, 2020). In contrast to TFR1, TFR2 has lower affinity to holo-Tf and is predominantly expressed in liver and erythroid precursor cells. Mutations of *TFR2* in human and germline or hepatocyte-specific deletion of *Tfr2* in mice cause type 3 hemochromatosis (Camaschella et al, 2000; Fleming et al, 2002). More recently, Tfr2 has been reported to regulate osteoblastic bone formation and bone mass in mice, independent of its function in iron homeostasis (Rauner et al, 2019). Nevertheless, the cell autonomous functions of TFR1 and TFR2 in osteoclast lineage cells remain unknown.

To elucidate the role of Tfr1 and Tfr1-mediated iron uptake in osteoclast lineage cells, we generated *Tfr1* conditional knockout mice in myeloid osteoclast precursors and mature osteoclasts by crossing *Tfr1*-floxed mice with *LysM*-Cre and *Ctsk*-Cre mice, respectively, and used them for comprehensive skeletal phenotyping and mechanistic studies.

## Results

### Tfr1-mediated iron uptake is the major route for iron acquiring in murine osteoclasts

It has been reported that Tf stimulates osteoclastogenesis whereas iron chelation inhibits this process (Ishii et al, 2009). In this study, we first set out to determine which of the four cellular iron uptake pathways in mammalian cells (i.e. Tf-dependent, heme, ferritin, and NTBI) function in osteoclasts. For this purpose, we quantified the mRNA level of genes encoding the major transporters along each iron acquiring pathway (supplemental Figure 1B) by real-time PCR in three different stages of murine osteoclast lineage cells: bone marrow monocytes, mononuclear osteoclast precursors, and multinucleated mature osteoclasts. By this assay, we observed that genes participated in Tf/Tfr-mediated iron uptake: *Tfrc* (encoding Tfr1), *Tfr2, Slc11a2* (encoding Dmt1) were upregulated in mature osteoclasts (Figure 1A). We have previously reported that Steap4, a ferrireductase with a critical role in the Tf-dependent pathway, is highly expressed in mature osteoclasts and plays an important role in cellular iron homeostasis in osteoclasts *in vitro* (Zhou et al, 2013). Together, all key mediators of Tf-dependent iron uptake pathway are up-regulated during osteoclast differentiation. By contrast, the mRNA expression of *Slc39a14* (encoding Zip14, a transporter mediating cellular iron entrance via NTBI) was decreased during osteoclast differentiation. The expression of *Slc46a1*, which encodes heme transporter Hcp1, was slightly increased in mature osteoclasts and the mRNA level of *Flvcr2*, which encodes another heme transporter, was low (Figure 1A). The mRNAs of *Timd2* and *Scara5*, which encode two plasma membrane transporters of ferritin, were undetectable in osteoclast lineage cells (data not shown).

Although both Tfr1 and Tfr2 mRNA expression increase during osteoclastogenesis, the level of *Tfr2 mRNA* was much lower than that of *Tfrc* (Figure 1A). Moreover, deletion of Tfr1 completely eliminated the uptake of Tf-^59^Fe in mature osteoclasts, indicating that Tfr1 is the predominant receptor for Tf-uptake and its deficiency is not compensated by Tfr2 in osteoclasts (left panel in Figure 1B). There was a dramatic increase in cellular iron level in mature osteoclasts compared to monocytes and pre-osteoclasts. Loss of Tfr1 in osteoclasts led to more than 50% decrease in total intracellular iron content (right panel in Figure 1B), suggesting that Tf-dependent iron uptake is a major route for iron acquirement in mature osteoclasts. We and others have previously found that the expression of Fpn, the only iron exporter identified so far in mammalian cells, decreases during osteoclast differentiation (Gu et al, 2015; Wang et al, 2018) and loss of Fpn in myeloid osteoclast precursors increases cellular iron and stimulates osteoclastogenesis *in vitro* and *in vivo* (Wang et al, 2018). Therefore, by transcriptional upregulation of Tf-dependent iron uptake pathway and concomitant downregulation of iron exporter Fpn, osteoclasts augment their cellular iron level to meet high energy demand during osteoclast differentiation and bone resorption. Nevertheless, the role of Tfr1 and Tf-dependent iron uptake in osteoclast biology and bone remodeling has not been elucidated using genetically modified mouse models.

### Loss of Tfr1 in myeloid osteoclast precursors increases trabecular bone mass of long bones in female mice

The homozygous *Tfr1* germline knockout mice are embryonically lethal (Levy et al, 1999). To elucidate the role of Tfr1 in osteoclast differentiation and bone resorption during postnatal bone modeling and remodeling, we generated *Tfr1* myeloid-specific conditional knockout mice on C57BL6 background by crossing *Tfr1*-floxed mice, in which the exons 3-6 of murine *Tfrc* gene were flanked by two loxP sites (Chen et al, 2015), with *LysM*-Cre mice. Our pilot study demonstrated that the single allele of *LysM-Cre* (*Tfr1*^flox/flox^;*LysM*^Cre/+^) only led to partial deletion of Tfr1 in osteoclast lineage cells whereas two copies of *LysM-Cre* (*Tfr1*^flox/flox^;*LysM*^Cre/Cre^ ) completely eliminated Tfr1 expression in osteoclasts (data not shown). Moreover, trabecular bone mass and structure in *Tfr1 Tfr1*^flox/flox^;*LysM*^Cre/+^ mice evaluated by µCT were indistinguishable from their littermate control mice (supplemental Figure 2). Therefore, we used *Tfr1*^flox/flox^;+/+ and *Tfr1*^flox/flox^;*LysM*^Cre/Cre^ mice as control and *Tfr1* myeloid conditional knockout (*Tfr1*^ΔlysM^) mice, respectively, for further *in vivo* and *in vitro* studies.

**Figure 2.**
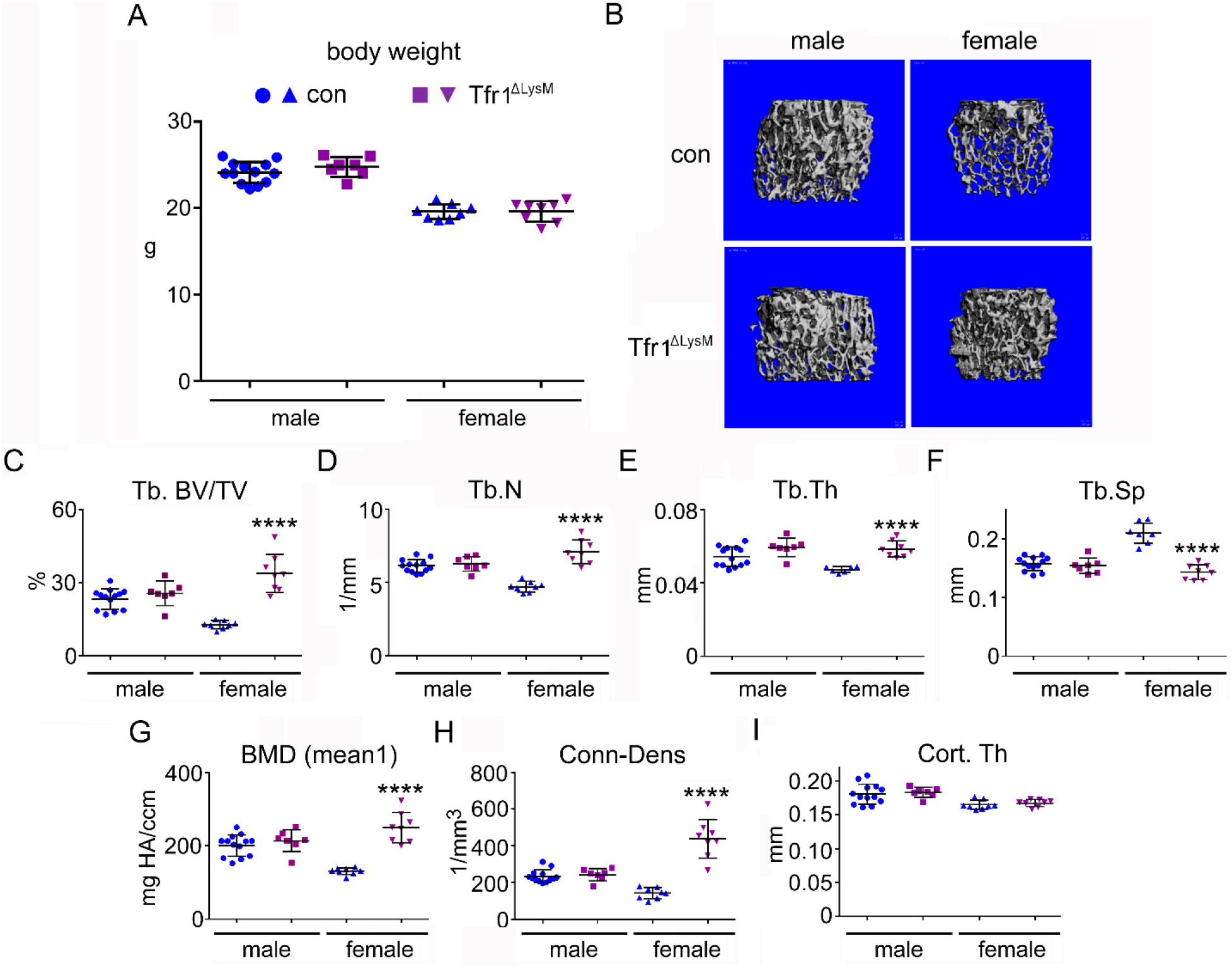
*Tfr1* myeloid conditional knockout mice develop normally and display increased trabecular bone mass in femurs of 10-week old female mice. (**A**) Body weight of 10-week old male and female control (con) and conditional knockout (Tfr1^ΔLysM^) mice in C57BL6J background. (**B**) Representative µCT images of distal femurs of male and female con and Tfr1^ΔLysM^ mice. (**C**) – (**I**) µCT analysis of trabecular and cortical bone mass and structure of distal femurs. Tb, trabecular bone; BV/TV, bone volume/tissue volume; Tb.N, trabecular number; Tb. Th, trabecular thickness; Tb.Sp, trabecular spacing; BMD, bone mineral density; Conn-Dens, connective density; Cort, cortical bone. *** p < 0.001; **** p < 0.0001 vs con by one-way ANOVA. n = 7-13.

Both male and female *Tfr1*^ΔlysM^ mice were born at the expected Mendelian ratio and developed normally with similar size and body weight compared to their littermate controls at 10 weeks old of age (Figure 2A). The µCT analysis of femurs revealed 2-fold more trabecular bone mass (BV/TV) with significant increase in trabecular number (Tb.N), trabecular thickness (Tb.Th), bone mineral density (BMD), connective density (Conn-Dens), and the decreased trabecular spacing (Tb.Sp) in femurs of female *Tfr1*^ΔlysM^ mice compared to controls. There was no difference in any of trabecular bone parameters in the male *Tfr1*^ΔlysM^ mice compared to their gender-matched controls (Figure 2B to 2H). Loss of Tfr1 in both male and female mice at this age had no effects on femoral cortical bone thickness (Figure 2I). The µCT analysis of tibias showed similar trabecular bone mass increase in female but not male *Tfr1*^ΔlysM^ mice (data not shown). Unexpectedly, this increased trabecular bone mass in the long bones of female *Tfr1*^ΔlysM^ mice was not observed in vertebral bones of 10-week old mice (supplemental Figure 3). Taken together, these results indicate that *Tfr1*-deletion in osteoclast precursors leads to increased trabecular bone mass and density in long bones of female mice.

**Figure 3.**
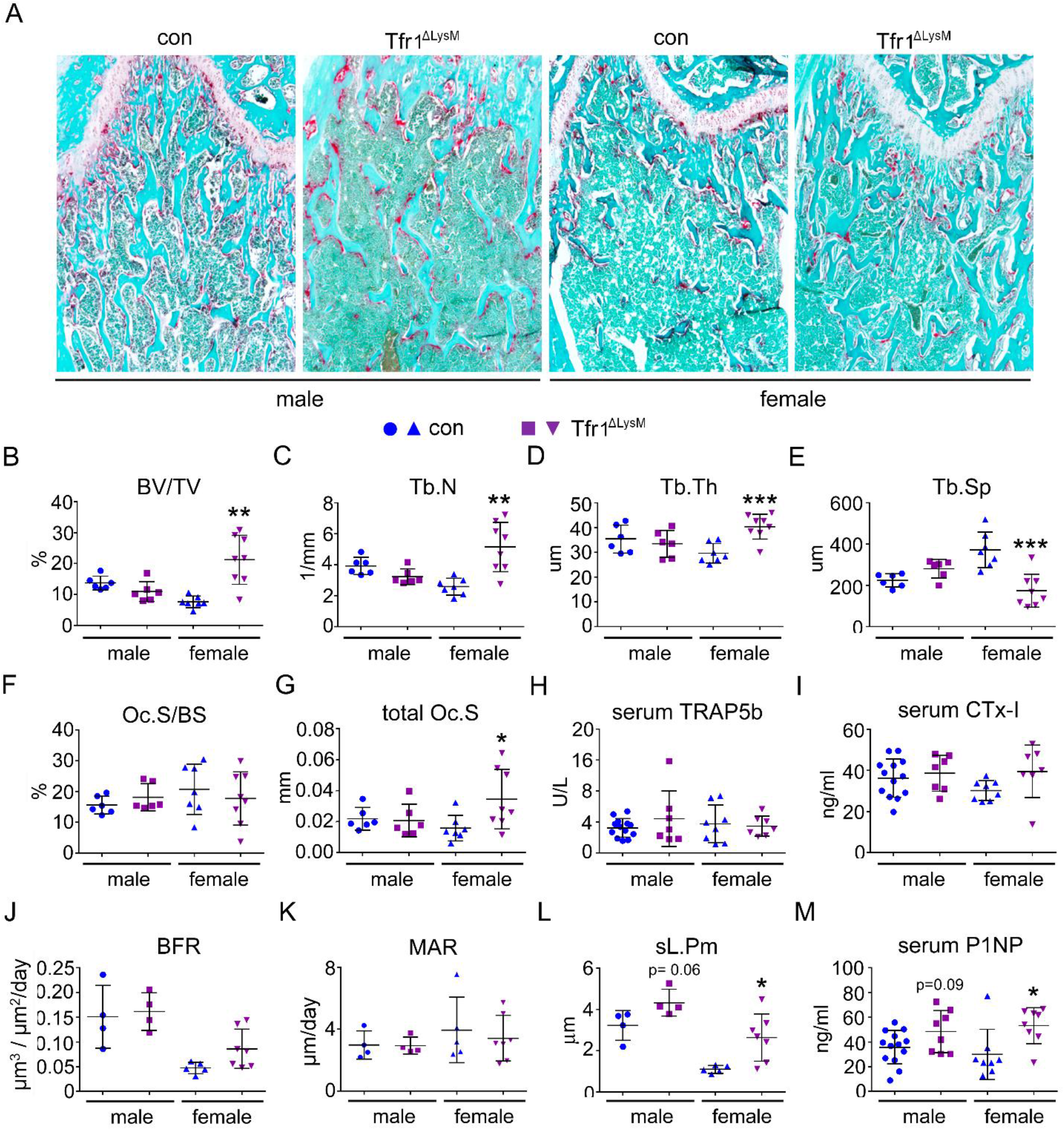
Loss of *Tfr1* in myeloid osteoclast precursors has no impacts on osteoclast number and bone formation in mice. (**A**) Images of Fast green and TRAP staining (10 × objective) and (**B**) – (**G**) Histomorphometric analysis of the metaphysis of decalcified distal femur histological sections of 10-week old male and female control (con) and Tfr1^ΔLysM^ mice. (**J**) – (**L**) Dynamic histomorphometry analysis of tetracycline-labeled sections from undecalcified distal femurs. (**H**), (**I**), and (**M**) quantitative measurements of serum markers for bone resorption and bone formation by ELISA. Tb, trabecular bone; BV/TV, bone volume/tissue volume; Tb.N, trabecular number; Tb. Th, trabecular thickness; Tb.Sp, trabecular spacing; OC.S/BS, osteoclast surface/bone surface; BFR, bone formation rate; MAR, mineral apposition rate; sL.Pm, single tetracycline labeled surface. * p < 0.05, ** p < 0.01, *** p < 0.001 vs con by one-way ANOVA. n = 4-13.

To determine the cellular processes that contribute to increased trabecular bone mass caused by deletion of *Tfr1* in myeloid cells, we performed histology and histomorphometry analysis of distal femurs from 10-week old male and female *Tfr1*^ΔlysM^ mice and their corresponding controls. Consistent with the µCT data, fast green and TRAP-staining of decalcified paraffin-embedded sections (Figure 3A) demonstrated increased trabecular mass, number, thickness, and decreased trabecular spacing in female but not male *Tfr1*^ΔlysM^ mice compared to their control littermates (Figure 3B to 3E). While the TRAP stained osteoclast surface adjusted for bone surface (Oc.S/BS), a histological parameter for osteoclast density, was the same in *Tfr1*^ΔlysM^ mice of either genders as in control mice (Figure 3F), the total Oc.S which reflects the number of osteoclasts was increased in female *Tfr1*^ΔlysM^ mice due to expansion of trabecular bone mass in these mice (Figure 3G). The systemic bone resorption level as measured by serum levels of bone resorption markers, TRAP5b and CTx-I, were similar in all genotypes of male and female mice (Figure 3H and 3I). Due to the increased total number of osteoclasts in female *Tfr1*^ΔlysM^ mice, the function of individual osteoclasts in female *Tfr1*^ΔlysM^ mice was decreased *in vivo*.

Since osteoclast-mediated bone resorption and osteoblast-mediated bone formation are coupled and regulate each other, we also examined bone formation activities in control and *Tfr1*^ΔlysM^ mice by dynamic bone histomorphometry analysis of tetracycline-labeled plastic-embedded tissue sections. As shown in Figure 3J and 3K, bone formation rate (BFR) and osteoblast mineral apposition rate (MAR) in trabecular bone were similar between control and *Tfr1*^ΔlysM^ male and female mice. The total tetracycline-labeled surface (sL.Pm) in female *Tfr1*^ΔlysM^ mice was higher than their control mice, likely due to increased trabecular bone mass/surface (Figure 3L). The increased serum level of P1NP (Figure 3M), a systemic bone formation marker, in female knockout mice may be caused by increased trabecular bone and total number of osteoblasts in these mice because the differentiation and bone formation activity of *Tfr1*-dificient osteoblasts cultured *in vitro* were the same as control cells (data not shown).

### Deletion of Tfr1 in mature osteoclasts increases trabecular bone mass of long bones in both male and female mice

To determine the role of Tfr1 in mature osteoclasts during bone remodeling, we crossed *Tfr1*-flox mice with *Ctsk*-Cre knock-in mice, a Cre-driver that has been extensively used to disrupt genes of interest in mature osteoclasts (Fujiwara et al, 2016; Wang et al, 2018; Nakamura et al, 2007). The progeny of *Tfr1*^+/+^;*Ctsk*^Cre/+^ and *Tfr1*^flox/flox^;Ctsk^Cre/+^) congenic mice on a C57BL6/129 mixed background were used as control and osteoclast conditional knockout (*Tfr1*^ΔCTSK^) mice, respectively. Both male and female *Tfr1*^ΔCTSK^ mice were born and developed normally. The body size and weight of 10-week old male and female *Tfr1*^ΔCTSK^ mice were similar to their littermate control mice (Figure 4A). The µCT examination of femurs unveiled a more than two-fold increase of trabecular BV/TV with increased trabecular number, thickness, and decreased trabecular spacing in both male and female *Tfr1*^ΔCTSK^ mice compared to their littermate controls (Figure 4B to 4F). A slight increase in cortical bone thickness was only observed in female *Tfr1*^ΔCTSK^ mice (Figure 4G). Similar to *Tfr1*^ΔlysM^ mice, loss of *Tfr1* in mature osteoclasts had little effects on trabecular bone mass and structure in vertebral bones of male and female *Tfr1*^ΔCTSK^ mice (supplemental Figure 4).

**Figure 4.**
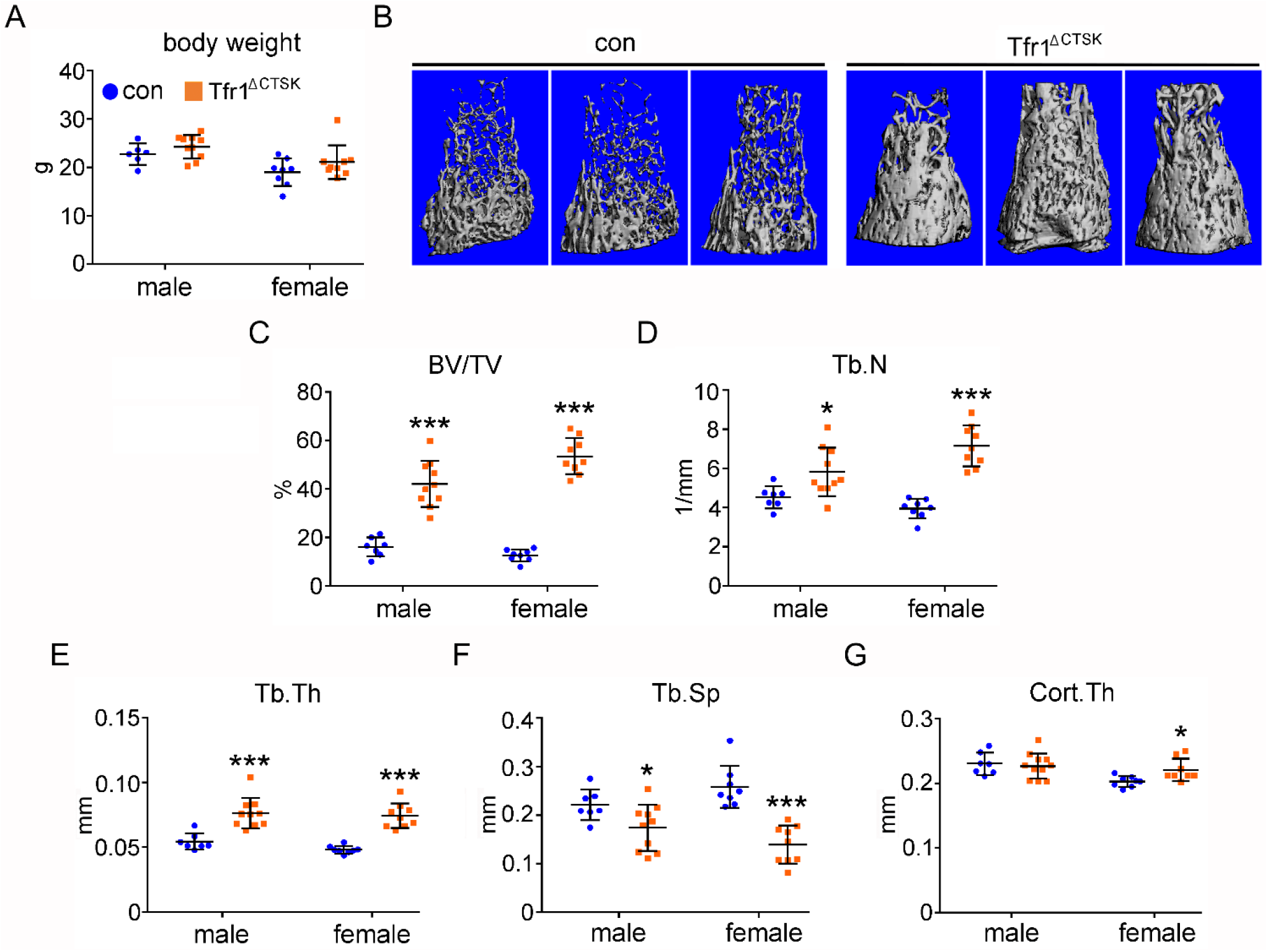
Deletion of *Tfr1* in cathepsin K-Cre expressing osteoclasts results in increased trabecular bone mass in distal femurs of 10-week old male and female mice. (**A**) Body weight of 10-week old male and female control (con) and conditional knockout (Tfr1^ΔCTSK^) mice in 129 × C57BL6J mixed background. (**B**) Representative µCT images of distal femurs of male and female con and Tfr1^ΔCTSK^ mice. (**C**) – (**G**) µCT analysis of trabecular and cortical bone mass and structure of distal femurs. Tb, trabecular bone; BV/TV, bone volume/tissue volume; Tb.N, trabecular number; Tb. Th, trabecular thickness; Tb.Sp, trabecular spacing; Cort, cortical bone. * p < 0.05, *** p < 0.001 vs con by one-way ANOVA. n = 7-10.

Histology and histomorphometry analysis of paraffin-embedded femoral sections demonstrated dramatic increase in trabecular BV/TV in both male and female *Tfr1*^ΔCTSK^ mice relative to their control mice (Figure 5A and 5B). These data are consistent with the µCT results shown in Figure 4. The number of TRAP-positive osteoclasts per bone perimeter (N.OC/Pm) as well as the percentage of osteoclast-covered bone surface (OC.Pm) in *Tfr1*^ΔCTSK^ mice of either genders were not different from control mice (Figure 5C and 5D). These results indicate that loss of *Tfr1* in Ctsk-Cre expressing osteoclasts increases trabecular bone mass but has no effects on osteoclastogenesis *in vivo*.

**Figure 5.**
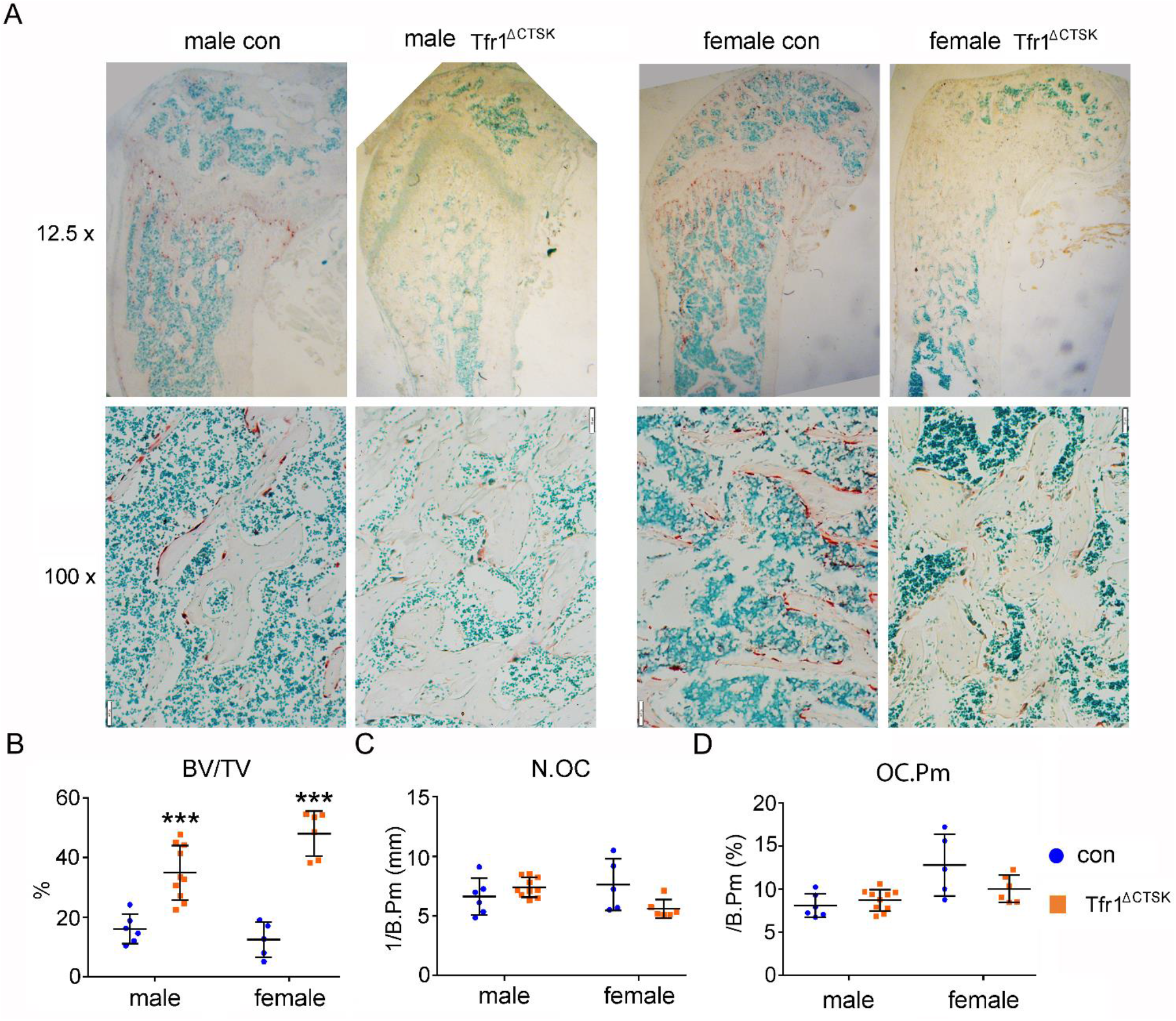
Deletion of *Tfr1* in late stage of osteoclast lineage cells increases trabecular bone mass without influence on osteoclastogenesis *in vivo*. (**A**) The low (12.5 ×) and high (100 ×) magnification of images of TRAP staining of histological sections of decalcified distal femurs of 10-week old control (con) and *Tfr1* conditional knockout (Tfr1^ΔCTSK^) mice. (**B**) – (**D**) Histomorphometric analysis of trabecular bone mass and osteoclast number. BV/TV, Trabecular bone volume/tissue volume; N.OC, osteoclast number/mm bone surface; OC.Pm, percentage of osteoclast surface/bone surface. *** p < 0.001 vs control by one-way ANOVA. n = 5-10.

### Tfr1-mediated iron uptake is dispensable for osteoclastogenesis but plays an important role in mature osteoclast actin cytoskeletal organization and bone resorption in vitro

To further identify the cellular and molecular mechanisms by which Tfr1 and Tfr1-mediated iron uptake regulate osteoclasts, we isolated bone marrow cells from C57BL6 male and female control and *Tfr1*^ΔlysM^ mice and cultured them with either M-CSF alone to generate bone marrow monocytes or M-CSF plus RANKL to induce mononuclear osteoclast precursor cells and multinucleated mature osteoclasts *in vitro*. As shown in Figure 6A, the complete deletion of Tfr1 by two copies of *LysM*-Cre in both male and female osteoclast lineage cells had little effects on the protein level of osteoclast transcription factor Nfatc1 and the induction of osteoclast acidic hydrolase Ctsk. Accordingly, the number of TRAP^+^ multinucleated osteoclasts was not different in *Tfr1*^ΔlysM^ cultures of either genders compared to their respective controls. However, the Tfr1- deficient osteoclasts displayed dramatic defect in mature osteoclast spreading (Figure 6B and 6C). Since actin cytoskeleton organization plays an essential role in osteoclast spreading and the formation of podosome-belts in osteoclasts cultured on plastic dishes and actin-rings in osteoclasts cultured on bone matrix, we next stained actin filaments with Alexa-488 conjugated Phalloidin in control and Tfr1-deficient male and female osteoclasts cultured on plastic and bone. While the podosome-belts and actin-rings formed normally in control osteoclasts, these actin-based structures were dysregulated in Tfr1-deficient male and female osteoclasts (Figure 7A-7D). Since formation of actin-rings is a hallmark of osteoclast activation and function, these results indicate that loss of Tfr1 in osteoclast suppresses osteoclast function. Corroborating this hypothesis, we found that Tfr1-deficient osteoclasts resorbed less bone than controls as revealed by the resorption-pit staining on cortical bovine bone slices (Figure 7E and 7F).

**Figure 6.**
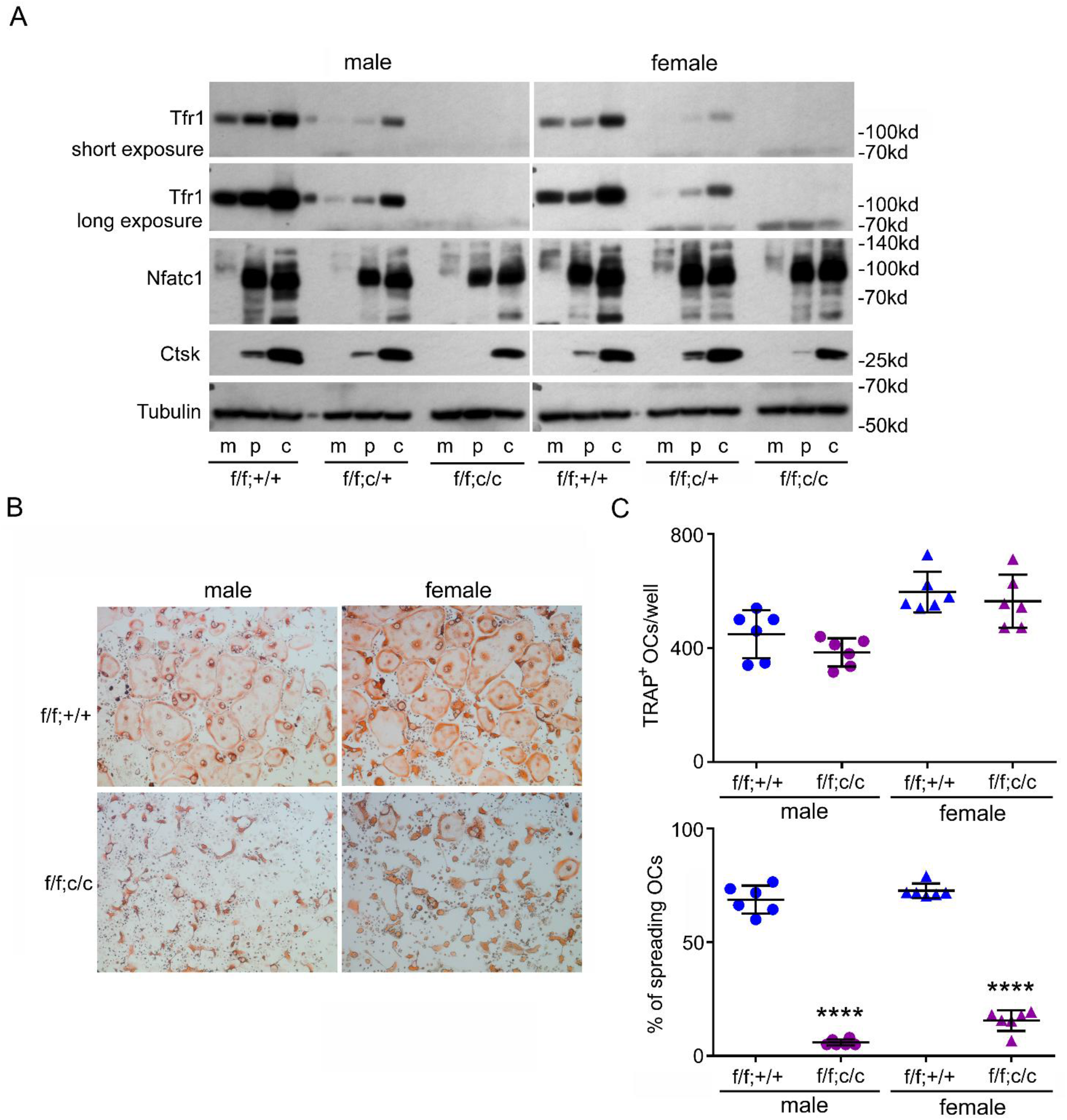
The complete deletion of Tfr1 by two alleles of LysM-Cre has no effects on osteoclast differentiation but induces spreading defect in mature osteoclasts. (**A**) The protein level of Tfr1 and osteoclast markers, Nfatc1and cathepsin K (Ctsk), in bone marrow monocytes (m), mono-nuclear pre- osteoclasts (p), and mature osteoclasts (c) was detected by western blotting. Tubulin served as loading control. (**B**) and (**C**) TRAP staining (4 × objective) and quantification of numbers of total and spreading osteoclasts (OCs). **** p < 0.0001 vs control (f/f;+/+) by one-way ANOVA. n = 6.

**Figure 7.**
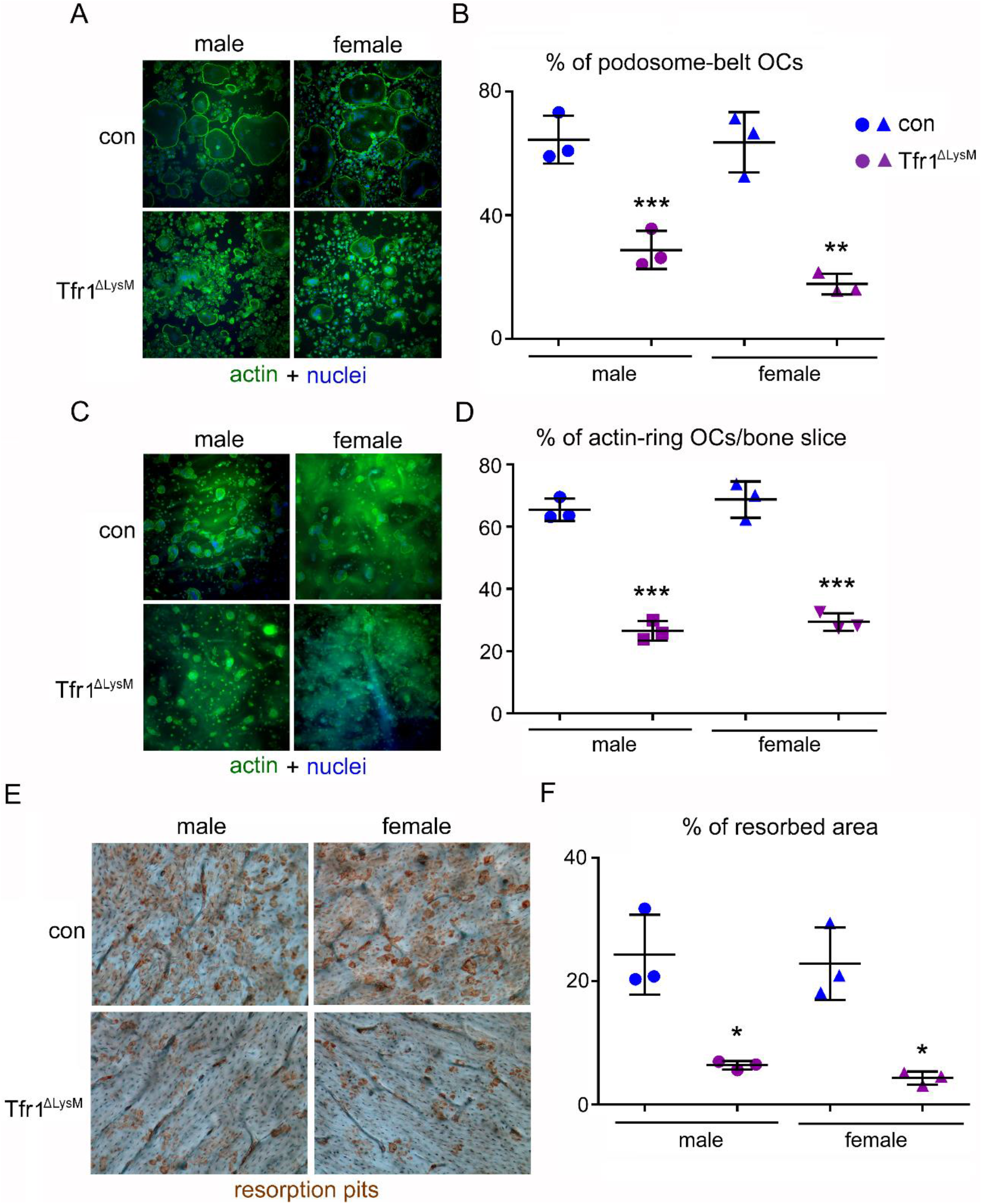
Loss of Tfr1 in osteoclast lineage cells attenuates actin cytoskeleton organization and inhibits bone resorption. (**A**) The actin filaments and nuclear were stained by Alexa-488 conjugated Phalloidin and Hoechst 33258, respectively, in control (con) and Tfr1^ΔLysM^ osteoclasts cultured on glass coverslips (20 × objective). (**B**) Quantification of the number of osteoclasts with podosome-belt from (**A**). (**C**) The actin filaments and nuclear were stained by Alexa-488 conjugated Phalloidin and Hoechst 33342, respectively, in osteoclasts cultured on cortical bovine bone slices (20 × objective). (**D**) Quantification of the number of osteoclasts with actin-rings from (**C**). (**E**) Resorption pits were stained by horseradish peroxidase conjugated wheat-germ agglutinin (10 × objective). (**F**) Quantification of resorbed area from (E). * p < 0.05, ** p < 0.01, *** p < 0.001 vs con by one-way ANOVA. n = 3.

Besides regulating iron-uptake, Tfr1 also plays noncanonical roles in regulation of intestinal epithelial homeostasis and mitochondrial morphology and function (Chen et al, 2015; Senyilmaz et al, 2015). To determine whether the defects in cytoskeleton organization and resorptive function of *Tfr1*-deficient osteoclasts were caused by lower intracellular iron content, we treated control and *Tfr1*^ΔlysM^ osteoclast cultures with increased doses of hemin and ferric ammonium citrate (FAC), which are transported into cells via heme transporters Hcp1 (Slc46a1) and NTBI transporter Zip14 (Slc39a14), respectively (supplemental Figure 1). While 10µM of hemin slightly stimulated osteoclast spreading, cytoskeleton organization, and bone resorption in control osteoclasts, hemin dose-dependently increased number of spreading osteoclasts in *Tfr1*-deficient osteoclasts (Figure 8A). In contrast, FAC had no effects on control or Tfr1-deficient osteoclasts (Figure 8B). At 10µM concentration, hemin significantly rescued the phenotypes of *Tfr1*-depleted osteoclasts (Figure 8C to 8E) Together with the data of mRNA expression of Hcp1 and Zip14 (Figure 1), these results suggest that heme but not NTBI and ferritin pathways is an alternative iron acquiring mechanism in osteoclasts in addition to Tfr1. Therefore, both our *in vivo* and *in vitro* findings unveil a critical role of Tfr1-mediated cellular iron homeostasis in osteoclast activation/function.

**Figure 8.**
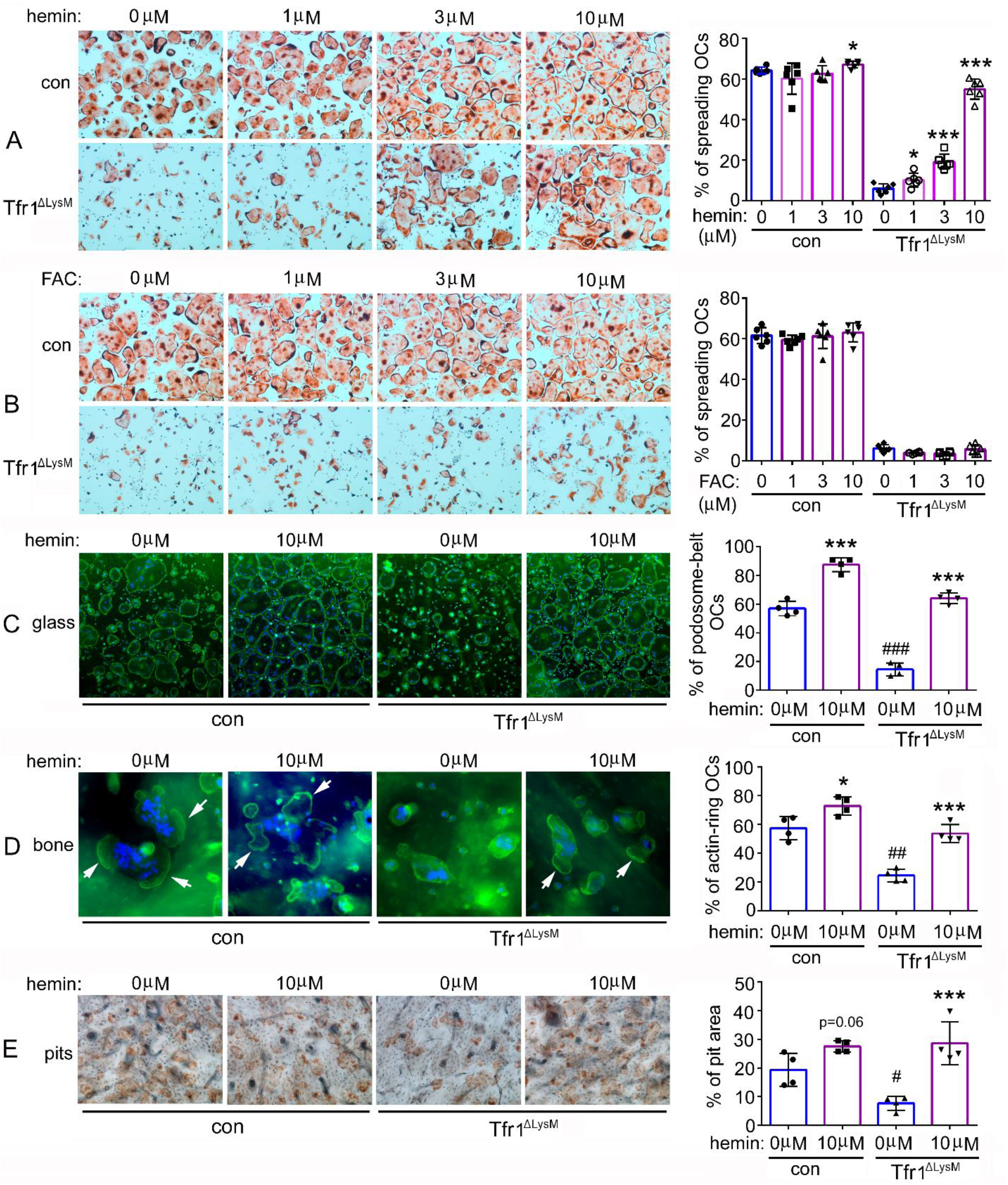
High dose of Hemin but not ferric ammonium citrate rescues the phenotypes of Tfr1-deficient osteoclasts. (**A**) and (**B**) TRAP staining and quantification of the number of spreading osteoclasts in control (con) and Tfr1^ΔLysM^ osteoclast cultures. n = 6. (**C**) and (**D**) Staining of actin filaments and nuclear and quantification of the number of podosome-belt and actin-ring osteoclasts cultured on glass coverslips and bone slices, respectively. n = 4. (**E**) resorption pit staining in con and cKO cultures. n = 4. * p < 0.05, *** p < 0.001 vs 0 µM control; # p < 0.05, ## p < 0.01, ### p < 0.001 vs con by one-way ANOVA.

### Depletion of Tfr1 in osteoclast lineage cells attenuates mitochondrial biogenesis, ROS production, and OXPHOS predominantly in mature osteoclasts

To determine which intracellular pathways are most affected by loss of Tfr1 in osteoclast lineage cells, we did quantitative proteomic analysis to identify proteins that are up- and down-regulated in Tfr1-deficient bone marrow monocytes, mononuclear precursors, and multinucleated mature osteoclasts. The heatmaps were generated using the ‘heatmap’ function in the R-package (Figure 9A). By a 1.2-fold cut-off, we found that a total of 135, 497, and 2562 proteins were influenced by loss of Tfr1 in monocytes, pre-osteoclasts, and mature osteoclasts, respectively (supplemental Tables 1 to 3). The abundance of proteins involved in Tf-dependent iron uptake pathway, including Tfr1, Steap3/4, DMT1 (Slc11a1), were all decreased in Tfr1-deficient cells (supplemental Table 1 to 3). The greater number of affected proteins in mature osteoclasts correlated with a high level of Tfr1 and Tfr1-mediated iron uptake in osteoclasts (Figure 1). By Ingenuity Pathway Analysis (IPA) analysis, we identified that the mitochondrial OXPHOS pathway and sirtuin signaling pathway were the most significantly down-and up-regulated pathways in mature osteoclasts, respectively (Figure 9B and supplemental Table 4). Among the mitochondrial proteins affected by loss of Tfr1 and Tf-dependent iron uptake in osteoclasts, the levels of proteins functioning in mitochondrial respiratory complex I to complex III decreased whereas the proteins participating in complex V, that lack heme and Fe-S clusters (supplemental Figure 1A), were increased probably by a compensate mechanism. The proteins in the complex IV were not affected by iron-deficiency (Figure 9C and supplemental Table 4).

**Figure 9.**
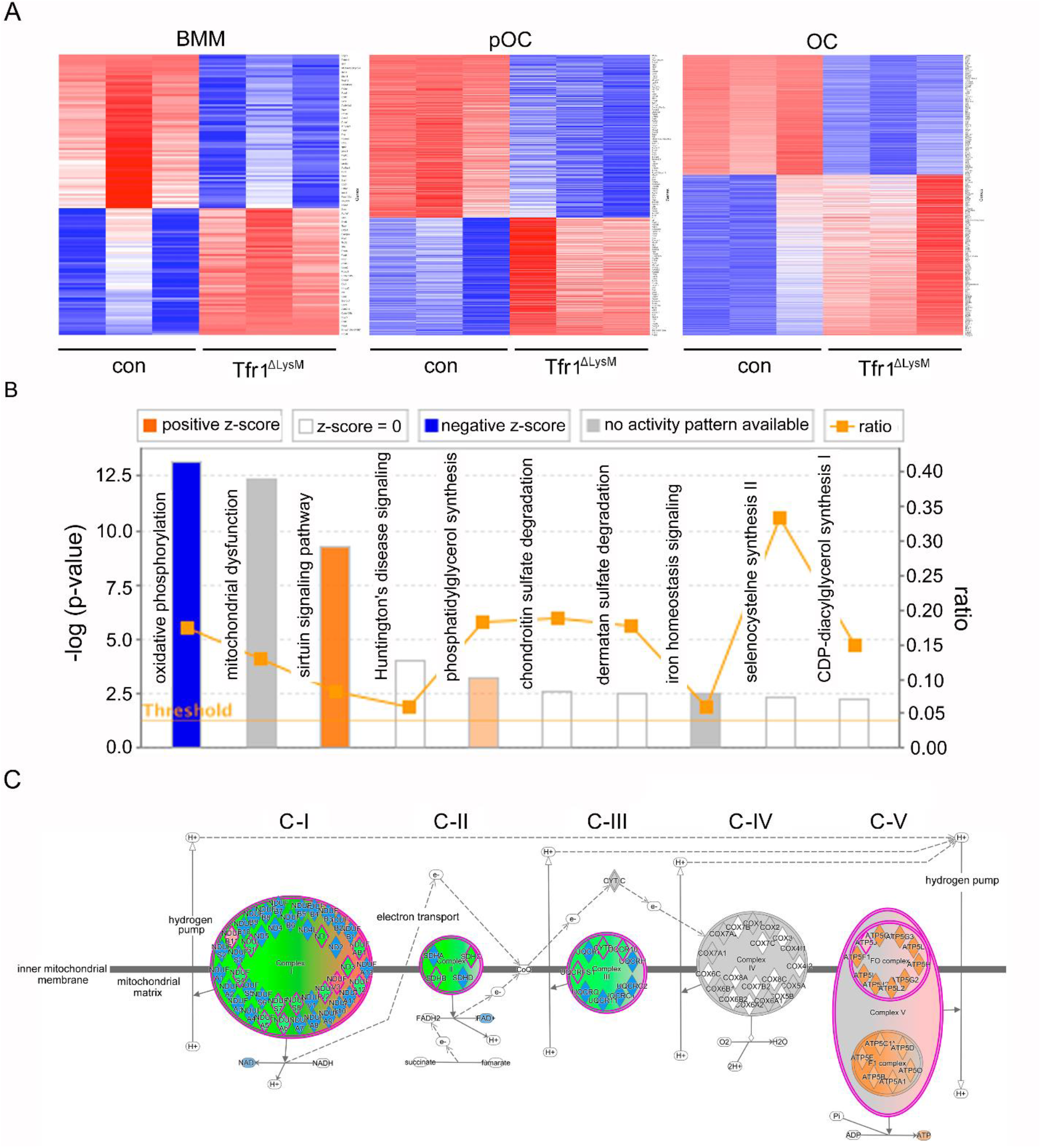
The mitochondrial oxidative phosphorylation is mostly affected by Tfr1-deficiency in mature osteoclasts. (**A**) Heat-maps of proteins that are differentially regulated in control (con) and Tfr1^ΔLysM^ bone marrow monocytes (BMM), mono-nuclear pre-osteoclasts (pOC), and mature osteoclasts (OC) identified by quantitative proteomics. (**B**) The signaling pathways that are affected by Tfr1-deficiency in mature osteoclasts identified by Ingenuity Pathway Analysis (IPA). (**C**) The changes of proteins along the mitochondrial respiration chain that are regulated by Tfr1 in mature osteoclasts. C-I to C-V, mitochondrial respiratory complex-I to complex-V.

To further characterize how iron-deficiency modulates mitochondrial biogenesis and metabolism in osteoclast lineage cells, we first stained the control and *Tfr1*^ΔlysM^ osteoclast lineage cells with fluorescence-labelled Mito Tracker. Manual quantification of Mito Tracker fluorescent intensity of individual cells showed a small decrease in mitochondrial mass in Tfr1-deficient mature osteoclasts but not in monocytes and pre-osteoclasts (Figure 10A). Next, we examined the mitochondrion-derived ROS by MitoSOX-staining of control and *Tfr1*^ΔlysM^ cultured cells. As demonstrated in Figure 10B, loss of Tfr1significantly decreased mitochondrial ROS production in monocytes and osteoclasts which was more pronounced in mature OCs. Insufficient iron in *Tfr1*^ΔlysM^ osteoclasts also resulted in a mild decrease in mitochondrial membrane potential (Figure 10C).

**Figure 10.**
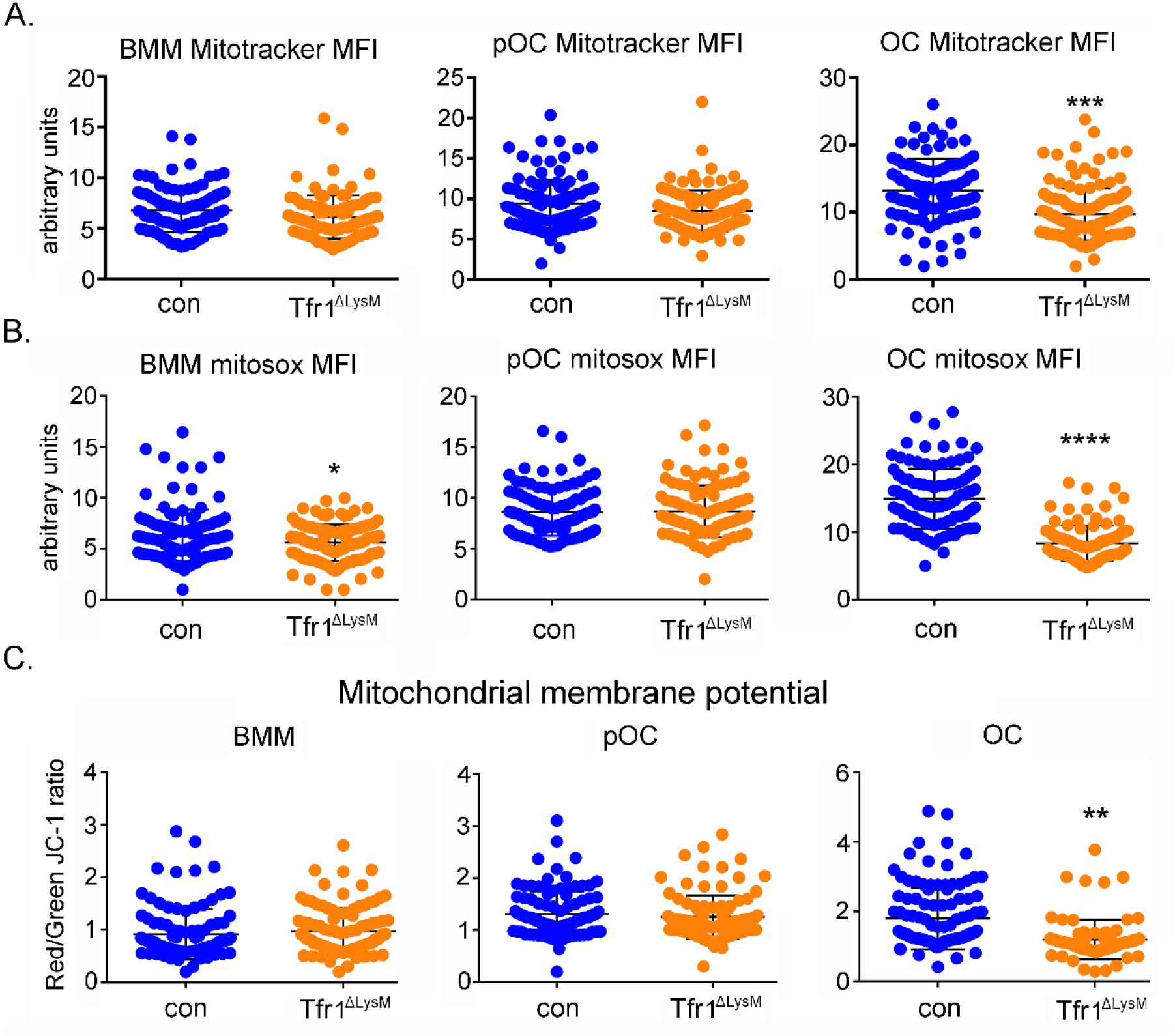
Loss of Tfr1 inhibits mitochondrial mass, ROS production, and membrane potential in mature osteoclasts. (**A**) Quantification of mitochondrial mass by MitoTracker Green staining in control (con) and Tfr1^ΔLysM^ bone marrow monocytes (BMM), mono-nuclear pre-osteoclasts (pOC), and mature osteoclasts (OC). (**B**) Measurement of mitochondria-derived ROS by Mitosox staining in con and cKO osteoclast lineage cells. (**C**) Measurement of mitochondrial membrane potential by JC-1 cationic carbocyanine dye staining in con and cKO osteoclast lineage cells. ** p < 0.01, *** p < 0.001, **** p < 0.0001 vs con by unpaired *Student’s t-test*. MFI, mean fluorescence intensity per cell in arbitrary units.

Lastly, we measured mitochondrial respiration using the Seahorse Extracellular Flux Analyzer in control and Tfr1-deficient osteoclast lineage cells isolated. As depicted in Figure 11A, the basal mitochondrial respiration is the direct measure of oxygen consumption rate (OCR) attributed to the mitochondrial electron transport chain (ETC). The maximum respiration is induced by incubating cells with the uncoupler, carbonyl cyanide 4-(trifluoromethoxy)phenylhydrazone (FCCP). The ATP-linked respiration is measured by exposing cells to oligomycin, an inhibitor of ATP synthase complex V. The nonmitochondrial respiration, which is the remaining OCR when the ETC activity is completely abolished by a mixture of rotenone/antimycin A. This parameter reflects the portion of cellular respiration by oxygen-consuming enzymes such as NADPH oxidases, heme oxygenases, and/or lipoxygenases. The residual OCR after inhibition by oligomycin is the measure of the protons pumped through ETC, which consumes oxygen without generating any ATP (due to inhibited ATP synthase activity by the drug) and is referred to the H^+^ leak. As shown in Figure 11B-11H, all these mitochondrial respiratory parameters increased dramatically in control and TfR1-null mature osteoclasts compared to their respective precursors. However, there were > 2-fold decreases of these measures in Tfr1-deficient mature osteoclasts relative to control osteoclasts. Furthermore, we calculated the reserve respiratory capacity, which is the difference between the maximum and basal respiration that can be utilized in the event of a sudden energy demand in cells. We reported this as a percentage for each group of cells by using the formula of [(maximum OCR - basal OCR)/maximum OCR × 100]. Again, the reserve respiratory capacity in Tfr1-null osteoclasts was greatly reduced when compared to control cells (Figure 11E). By contrast, there was no difference in the coupling efficiency (the ratio of ATP-linked OCR to basal OCR) of control and *Tfr1*^ΔlysM^ osteoclast lineage cells (Figure 11G).

**Figure 11.**
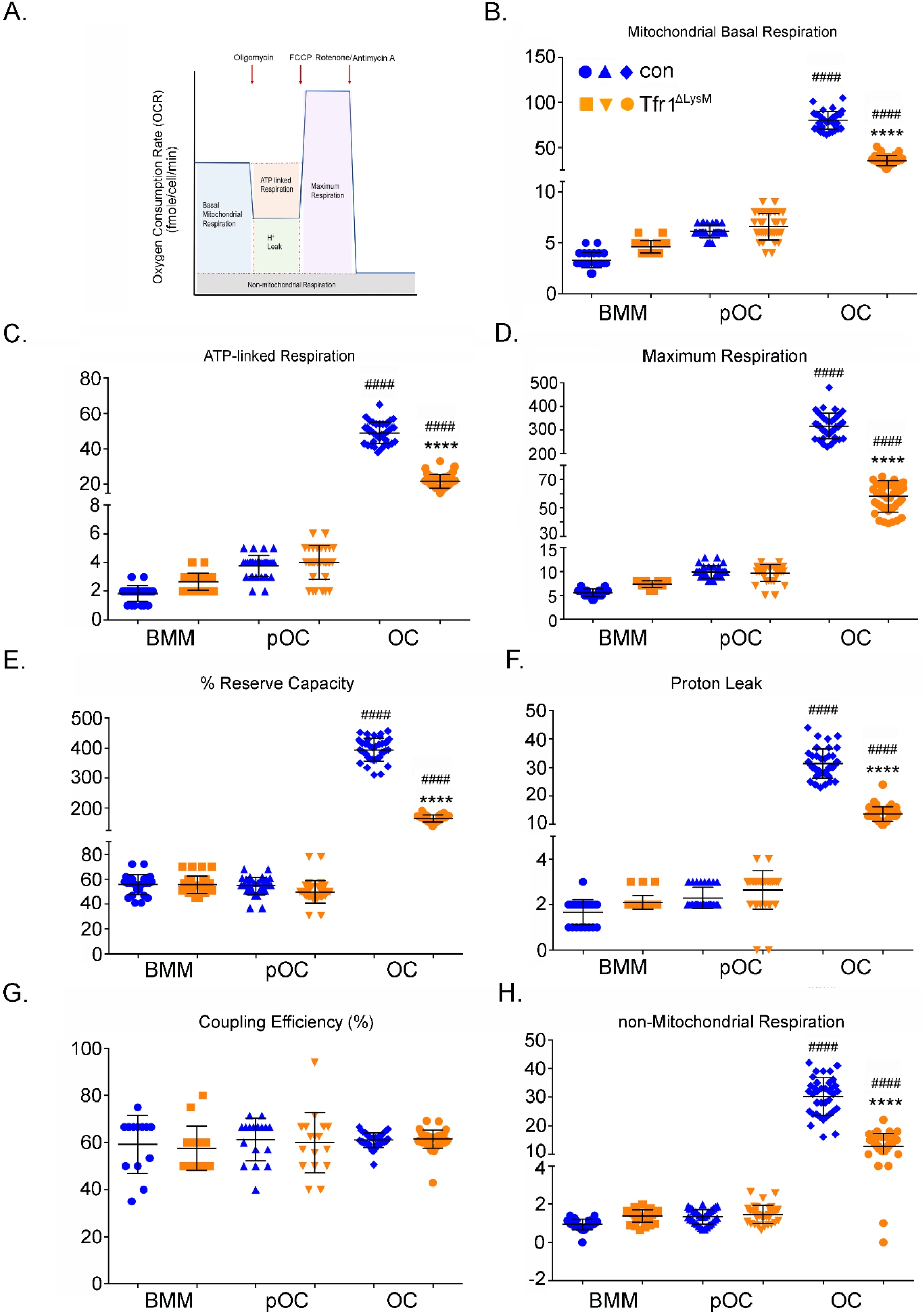
Tfr1-deletion in osteoclast lineage cells impairs mitochondrial and non-mitochondrial respirations in mature osteoclasts. (**A**) A graphic illustration of oxygen consumption measured by a Seahorse Extracellular Flux analyzer. FCCP, carbonyl cyanide p-trifluoro-methoxyphenyl hydrazone, a synthetic mitochondrial uncoupler. (**B**) – (**H**) Different fractions of mitochondrial and non-mitochondrial respirations per cell of control (blue) and Tfr1^ΔLysM^ (orange) bone marrow monocytes (BMM), mono-nuclear pre-osteoclasts (pOC), and mature osteoclasts (OC). #### p < 0.0001 vs BMM and pOC; **** p < 0.0001 vs control OC by unpaired *Student’s t-test*.

### TfR1-mediated iron uptake modulates osteoclast actin cytoskeleton through the WAVE regulatory complex

To identify the mechanisms by which cellular iron and energy metabolism regulate osteoclast cytoskeleton, we turned to analyze the level of key osteoclast cytoskeleton regulating proteins (Blangy et al, 2020) in Tfr1-deficient osteoclast lineage cells relative to their control cells in our quantitative proteomic data bases (supplemental Table 1-3). More cytoskeleton proteins in mature osteoclasts were affected by Tfr1-dificiency than those in osteoclast precursors (supplemental Table 5). Among them, 25 cytoskeleton proteins were down-regulated whereas 17 proteins were up-regulated by more than 1.2-fold in Tfr1-dificient osteoclasts compared to control cells. 15 osteoclast cytoskeleton-regulatory proteins remained the same between control and Tfr1-dificient osteoclasts. IPA analysis of the cytoskeleton-regulating pathways affected by Tfr1-deficiency in osteoclasts (Figure 12A) unveiled that the protein level of several molecules involved in activation of β_3_-integrin pathway, which is critical for osteoclast cytoskeleton organization and bone resorption (Teitelbaum, 2011), were decreased in Tfr1-null osteoclasts including β_3_-integrin (Itgb3), c-Src, Rap1a/b, Rap2b, Rapgef1, Rap1gds (supplemental Table 5). A few actin-bundling and regulating proteins such as myosin heavy chain 14 (Myh14), α-actinin 1 and 4 (Actn1 and Actn4), and filamin A (Flna) were also down-regulated in Tfr1-depleted osteoclasts. The branching and polymerization of filamentous actin in eukaryotic cells is mediated by the Arp2/3 complex which is controlled by the WAVE regulatory complex (WRC). Mammalian WRC contains five subunits of Cyfip1/2, Hem1/2 (encoded by *Nckap1l* and *Nckap1*, respectively), Abi1/2/3, HSPC300, and WAVE1/2/3 (Rottner et al, 2021). The WRC is activated by the small GTPases Rac1 and Arf1 through their direct binding to Cyfip1/2 and Hem1/2. Strikingly, 12 out of 25 down-regulated cytoskeletal proteins in Tfr1-null osteoclasts (supplemental Table 5) are the components of the Rac1/Arf-WRC-Arp2/3 axis. These data indicate that Tfr1-mediated iron uptake regulates osteoclast actin cytoskeleton organization, at least in part, through modulating the stability of the WRC-Arp2/3 actin-regulating module. To test this hypothesis, we constructed a recombinant retroviral vector expressing murine Hem1. After retroviral transduction in Tfr1- deleting bone marrow monocytes, we confirmed the expression of recombinant Hem1 in osteoclast lineage cells by western blotting (Figure 12B). The overexpression of Hem1 partially rescued the cytoskeleton organization defect in Tfr1-null osteoclasts as demonstrated by a significant increase in the number of well-spreading, podosome-belt containing osteoclasts compared to non-transduced and empty vector transduced Tfr1-null osteoclasts (Figure 12C).

**Figure 12.**
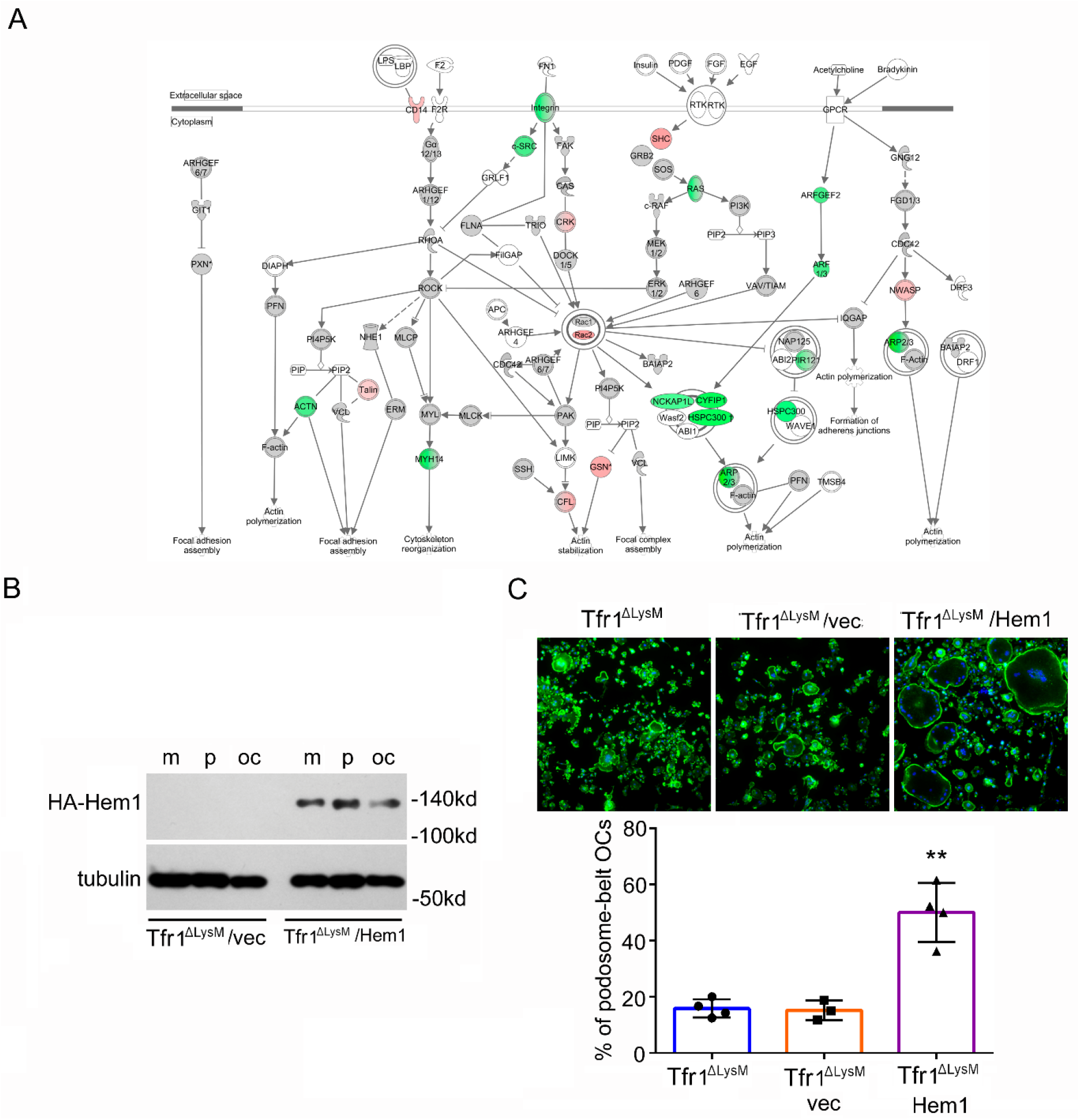
Loss of Tfr1 in osteoclasts reduces the level of cytoskeleton-regulating proteins and overexpression of Hem1 recuses the cytoskeletal organization defect in Tfr1-null osteoclasts. (A) A schematic map of cytoskeletal pathways generated by Ingenuity Pathway Analysis (IPA). The down-regulated proteins are shown in green and up- regulated proteins are marked in red orange. (B) Detection of HA-tagged Hem1 (encoded by Nckap1l) by western blotting in retroviral transduced Tfr1^ΔLysM^ bone marrow monocytes (m), mononuclear pre-osteoclasts (p), and mature osteoclasts (oc) expressing empty vector (vec) and recombinant Hem1. Tubulin served as loading control. (C) The staining of actin filaments and nuclear in osteoclasts cultured on glass coverslips and quantification of the number of osteoclasts with podosome-belt. ** p < 0.01 vs Tfr1^ΔLysM^ and Tfr1^ΔLysM^/vec by one-way ANOVA. n = 4.

## Discussion

Energy metabolism plays a pivotal role in osteoclast differentiation and function *in vitro* and *in vivo* (Kubatzky et al, 2018; Arnett and Orriss, 2018). Increased iron uptake coupled with the elevated expression of PGC1-β, a transcriptional coactivator of genes involved in energy metabolism, stimulate mitochondrial biogenesis and respiration to meet high energy demand during osteoclast differentiation and bone resorption activities (Ishii et al, 2009). Although Tf has been reported to promote osteoclastogenesis and bone resorption *in vitro* and iron chelation by desferrioxamine (DFO) has been shown to inhibit osteoclasts and prevent bone loss induced by estrogen deficiency in ovariectomized mice (Ishii et al, 2009), the precise functions of Tfr1 and Tfr1-mediated iron uptake in osteoclast biology and skeletal homeostasis remain incompletely understood. To fill in this knowledge gap, we have generated two lines of *Tfr1* conditional knockout mice in which Tfr1 expression is disrupted in myeloid osteoclast precursors and mature osteoclasts, respectively. Our *in vivo* skeletal phenotyping and *in vitro* mechanistic studies unveil a critical role of TfR1-mediated iron uptake in regulating osteoclast mitochondrial metabolism and cytoskeleton organization/bone resorption but not osteoclast differentiation. Our findings provide the experimental evidence that disruption of Tfr1 expression in osteoclast lineage cells in mice leads to a dramatic increase in trabecular bone mass in long bones of female mice.

The real-time qPCR detection of mRNA expression of transporters and regulators in four mammalian iron-acquiring pathways (Tf-dependent, heme, ferritins, and NTBI) in osteoclast precursor and mature cells demonstrated that the key molecules involved in Tf/TFR-mediated iron uptake are significantly up-regulated during osteoclast differentiation. The expression of heme transporter Hcp1 is also increased in mature osteoclasts whereas the level of NTBI iron transporter Zip14 and ferritin transporter (Scara-5 and Tim-2) is either decreased in osteoclasts or undetectable in osteoclast lineage cells (Figure 1). These results indicate that osteoclasts obtain extracellular iron largely through uptake of Tf and heme. This notion is further supported by the evidence that high dose of hemin but not FAC is able to rescue the Tfr1-deficient osteoclast phenotypes (Figure 8). Furthermore, loss of Tfr1 in osteoclasts completely abolishes the uptake of Tf while Tfr2- deletion in monocytes has no effects on iron homeostasis (Rishi et al, 2016), indicating that Tfr1 is the major receptor for Tf and Tf-dependent iron uptake in osteoclast lineage cells.

At post pubertal age of 10 weeks, loss of Tfr1 in osteoclast precursors causes a two to three-fold increase in trabecular bone parameters in female but not male mice (Figure 2 and Figure 3), although both male and female Tfr1-null osteoclasts exhibit similar defects *in vitro* (Figure 6 and Figure 7). In contrast, disruption of Tfr1 expression in cathepsin K-expressing mature osteoclasts causes significant increase of trabecular bone mass in both genders of mice at 10 weeks, albeit the phenotype is more pronounced in female knockout mice (Figure 4 and Figure 5). The exact mechanisms underlying the gender difference in the trabecular bone phenotype of mice with Tfr1- deficiency in osteoclasts are unknown. A similar finding has been recently reported in *Glut1* myeloid conditional knockout mice (Li et al, 2020). While deletion of Glut1 in LysM-expressing myeloid cells in mice inhibits osteoclastogenesis and leads to increased trabecular bone mass in female but not male mice, Glut1-depleted bone marrow monocytes cultured from both male and female knockout mice have similar degree of blunted osteoclast differentiation *in vitro*. Estrogen has been reported to regulate systemic and cellular iron homeostasis by inhibiting the expression of hepcidin, a liver-derived iron regulating hormone that binds to and induces degradation of iron exporter Fpn (Hou et al, 2012; Yang et al, 2012; Ikeda et al, 2012). Thus, estrogen may synergistically reduce cellular iron content with Tfr1-deficiency in osteoclast lineage cells by enhancing iron export through hepcidin/ferroportin axis *in vivo*. In addition, we have recently shown that estrogen attenuates mitochondrial OXPHOS and ATP production in precursors but not mature osteoclasts (Kim et al, 2020). Therefore, it is likely that estrogen and Tfr1-deletion additively inhibit mitochondria metabolism in osteoclast precursor cells. This may explain why the sexual dimorphism of skeletal phenotype is more obvious in *Tfr1* myeloid knockout mice than in *Tfr1* mature osteoclast knockout mice. Nevertheless, the effects of estrogen on Tfr1 expression and cellular iron homeostasis in osteoclast precursor and mature osteoclasts need further investigation. Furthermore, little is known about how androgen directly modulates Tfr1 and iron status *in vivo* and *in vitro*.

Up-regulation of glycolysis and mitochondrial OXPHOS bioenergetic pathways are required for osteoclast differentiation and function (Kubatzky et al, 2018; Arnett and Orriss et al, 2018). In this study, we have demonstrated that both mRNA and protein of Tfr1 are most abundant in mature osteoclasts (Figure 1 and Figure 6). Disruption of Tfr1 expression in myeloid osteoclast precursor cells attenuates mitochondria biogenesis and respiration in mature but not progenitor osteoclasts, leading to defective osteoclast cytoskeleton organization and impeded bone-resorbing activities with little influence on osteoclast differentiation *in vitro* and *in vivo* (Figure 6 to 12). These results indicate that energy metabolism regulated by Tf11-mediated iron uptake is specifically indispensable for osteoclast activation and function but not for osteoclastogenesis. In contrast, Ishii et al have reported that *in vitro* knock-down of Tfr1 expression in bone marrow monocytes by short-hairpin RNA (shRNA) inhibits osteoclast differentiation (Ishii et al, 2009). This discrepant finding may be caused by the off-target effects of Tfr1 shRNA on expression of genes essential for osteoclast differentiation or by the distinct effects of short-term Tfr1 down-regulation by shRNA and long-term genetic deletion of Tfr1 on osteoclast lineage cells.

It has been recently reported that deletion of mitochondria coactivator PGC-1β in myeloid osteoclast precursor cells by *LysM*-Cre diminishes mitochondrial biogenesis and function in osteoclasts, leading to cytoskeletal disorganization and bone resorption retardation with normal osteoclasts differentiation (Zhang et al, 2018). The PGC-1β-stimulated osteoclast cytoskeleton activation is mediated by GIT1 (G protein coupled receptor kinase 2 interacting protein 1), an upstream activator of small GTPases Rac1 and Cdc42 that are indispensable for osteoclast cytoskeletal organization and function (Jones and Katan, 2007; Menon et al, 2010). Another report by Nakano et al has shown that deletion of G-protein Gα_13_ in myeloid cells augments mitochondrial biogenesis and metabolism in osteoclasts. The gain-of-function of mitochondrial pathway induced by Gα_13_-deletion in osteoclasts promotes actin cytoskeleton dynamic and bone resorption via activation of cytoskeleton regulators c-Src, Pyk2, and RhoA-Rock2. Again, Gα_13_-deletion in osteoclast lineage cells has minimum effects on osteoclastogenesis *in vivo* and *in vitro* (Nakano et al, 2019). These reports are consistent with our findings in that energy metabolism regulated by Tfr1-mediate iron uptake is primary for osteoclast cytoskeleton organization rather than osteoclast differentiation. Intriguingly, Tfr1 regulates osteoclast cytoskeleton probably via a distinct mechanism. The level of GIT1, Rac1, Cdc42, and Rock2 remains unchanged in Tfr1-null osteoclasts (supplemental Table 5). However, the components of the actin-regulating complex WRC and the isoforms of small GTPases Arf family members are mostly down-regulated, indicating that Tfr1 modulates osteoclast cytoskeleton through promoting the stability of WRC complex. In supporting of this premise, overexpression of Hem1 (also known as Nckap1l) rescues the cytoskeletal defects in Tfr1-deficient osteoclasts (Figure 12).

In summary, we have provided evidence demonstrating that Tfr1-mediated iron uptake is a major iron acquisition pathway in osteoclast lineage cells that differentially regulates trabecular bone remodeling in perpendicular and axial bones of female and male mice. The increased intracellular iron facilitated by Tfr1 is specifically required for osteoclast mitochondrial energy metabolism and cytoskeletal organization.

## Materials and Methods

### Mice and genotyping

*Tfr1*-flox congenic mice on a 129Sv background were kindly provided by Dr. Nancy C Andrews (Duke University). *Tfr1*-flox mice on a homogeneous C57BL6 background were generated by backcrossing 129Sv *Tfr1*-flox mice with C57BL6 mice for more than 10 generations. *LysM*-Cre mice on a C57BL6 background (B6.129P2^tm1/cre^, stock number 004781) were purchased from The Jackson Laboratory (Bar Harbor, Maine, USA). *Ctsk-*Cre congenic mice on a C57BL6 background were obtained from Dr. Takashi Nakamura (Keio University, Tokyo, Japan). The primers and PCR protocols for genotyping *Tfr1*-flox and *LysM*-Cre mice followed those provided by The Jackson Laboratory. The following primers for genotyping *Ctsk-*Cre mice were used: P1N (5’-CCTAATTATTCCTTCCGCCAGGATG-3’), P2N (5’- CCAGGTTATGGGCAGAGATTTGCTT-3’), and P3N (5’- CACCGGCATCAACGTTTTCTTTTCG-3’). *In vivo* analyses of skeletal phenotypes were performed on F2 mice of mixed and homogeneous backgrounds.

### Reagents and antibodies

Alexa Fluor-488 Phalloidin (catalog no. A12379), alpha-MEM (catalog no. 78-5077EB), Hoechst 33342 (catalog no. H3570), and 10 × Trypsin/EDTA (catalog no. 15400-054) were purchased from Thermo-Fisher Scientific. cOmplete EDTA-free protease inhibitor cocktail (catalog no. 4693159001); 3,3’-diaminobenzidine (DAB) tablets (D-5905), High glucose DMEM (catalog no. D-5648), enhanced chemiluminescent detection reagents (ECL, catalog no. WBKLS0100), ferric ammonium citrate (FAC, catalog no. F5879), 30% H_2_O_2_ (catalog no. 216763), hemin (catalog no. H9039), NaK tartrate (catalog no. S6170), Napthol AS-BI phosphoric acid solution (catalog no. 1802), 10 × penicillin–streptomycin-l-glutamine (PSG) (catalog no. G1146), peroxidase-conjugated WGA (Wheat germ agglutinin) lectin (catalog no. L-7017), polyvinylidene difluoride membrane (PVDF, catalog no. IPVH00010), 1 × RIPA buffer (catalog no. R-0278), and mouse apo-transferrin (catalog no. T0523) were obtained from MilliporeSigma.

Fetal bovine serum (FBS) was purchased from Hyclone. Blasticidin (catalog no. 203350) was bought from EMD Chemicals/Millipore. TransIT-LT1 transfect reagent (catalog no. MIR2300) was obtained from Mirus Bio LLC. Fe^59^ in ferric chloride form was purchased from PerkinElmer Inc. All other chemical compounds were from MilliporeSigma.

The antibodies used in this study were obtained from the following resources: mouse monoclonal anti-cathepsin K (clone 182-12G5, catalog no. MAB3324, MilliporeSigma); mouse monoclonal anti-HA.11 antibody (clone 16B12, catalog no. 901513, Biolegend); mouse monoclonal anti-NFATc1 (catalog no. sc-7294, Santa Cruz Biotechnology); mouse monoclonal anti-transferrin receptor (clone H68.4 catalog no. 13-6800, Thermo-Fisher Scientific); mouse monoclonal anti-tubulin (clone DM1A, catalog no. T9026, MilliporeSigma); horseradish peroxidase conjugated anti-mouse secondary antibody (catalog no. 7074, Cell Signaling Technology); and horseradish peroxidase conjugated anti-rabbit secondary antibody (catalog no. 7076, Cell Signaling Technology).

### µCT

The left femurs, tibias, and L4 vertebrae were cleaned of soft tissues and fixed in 4% paraformaldehyde in PBS for overnight at 4°C. After washing with PBS for three times, the bones were stored in PBS with 0.02% sodium azide. The bones were loaded into a 12.3-mm diameter scanning tube and were imaged in a μCT (model μCT40, Scanco Medical).We integrated the scans into 3-D voxel images (1024 x 1024 pixel matrices for each individual planar stack) and used a Gaussian filter (sigma = 0.8, support = 1) to reduce signal noise. A threshold of 200 was applied to all scans, at medium resolution (E = 55kVp, I = 145μA, integration time = 200ms).

### Histology and bone histomorphometry

The left femurs were embedded undecalcified in methyl methacrylate. The dynamic histomorphometric examination of trabecular bone formation was done on 5μm longitudinal sections with a digitizer tablet (OsteoMetrics, Inc., Decatur, GA, USA) interfaced to a Zeiss Axioscope (Carl Zeiss, Thornwood, NY, USA) with a drawing tube attachment. The right femurs fixed and decalcified in 14% EDTA for 7-10 days. The bones were embedded in paraffin before obtaining 5-μm longitudinal sections. After removal of paraffin and rehydration, sections were stained for TRAP activity and counter-stained with fast green and osteoclasts were enumerated on the trabecular bone surface using Osteomeasure histomorphometric software (OsteoMetrics, Inc., Decatur, GA, USA).

### Serum TRAcP-5b, CTx-I, and PINP ELISA

Blood were collected retro-orbitally under inhalation of a 2% isoflurane/oxygen mix anesthesia immediately prior to sacrifice. Serum was obtained by centrifugation of blood in a MiniCollect tube (catalogue no. 450472, Greiner Bio-one GmbH, Austria). The serum levels of TRAcP-5b, CTx-I, and PINP were measured by a mouse TRAP (TRAcP 5b) kit (SB-TR103), RatLaps (CTx-I) EIA (AC-06F1), and rat/mouse PINP EIA kit (AC-33F1) from Immunodiagnostic Systems following their instructions.

### In vitro osteoclast cultures

Whole bone marrow was extracted from tibia and femurs of 6- to 8- week-old control and conditional knockout male and female mice. Red blood cells were lysed in buffer (150 mM NH4Cl, 10 mM KNCO3, 0.1 mM EDTA, pH 7.4) for 5 minutes at room temperature. 5 × 10^6^ bone marrow cells were plated onto a 100mm petri-dish and cultured in α-10 medium (α-MEM, 10% heat-inactivated FBS, 1 × PSG) containing 1/10 volume of CMG 14–12 (conditioned medium supernatant containing recombinant M-CSF at 1μg/ml) for 4 to 5 days. Bone marrow monocytes were lifted by 1 × Trypsin/EDTA and replated at density of 160/mm^2^ onto tissue culture plates or dishes with 1/100 vol of CMG 14–12 culture supernatant along for monocytes or cultured with 1/100 vol of CMG 14–12 culture supernatant plus 100 ng/ml of recombinant RANKL for 2 and 4 days to generate mononuclear pre-osteoclasts and mature osteoclasts, respectively.

### Retroviral transduction

A total of 3μg of pMX retroviral vector or recombinant pMX vector expressing HA-tagged murine Hem1 were transfected into Plat E retroviral packing cells using TransIT-LT1 transfection reagent. Virus supernatants were collected at 48 hours after transfection. 5 × 10^6^ bone marrow cells isolated from TfR1 myeloid conditional knockout mice were plated onto a 100mm petri-dish and cultured in α-10 medium (α-MEM, 10% heat-inactivated FBS, 1 × PSG) containing 1/10 volume of CMG 14–12 supernatant for two days. Bone marrow monocytes were transduced with viruses for 24 h in α-10 medium containing 1/10 volume of CMG 14–12 supernatant and 20μg/ml of protamine. Cells were then lifted by 1 × Trypsin/EDTA and replated at 2 × 10^6^ density onto a 100mm petri-dish. The positively transduced cells were selected in α-10 medium containing M-CSF and 1.5μg/ml of blasticidin (203350, EMD Chemicals) for 3 days.

### TRAP staining

Osteoclasts cultured on 48-well tissue culture plate were fixed with 4% paraformaldehyde/PBS for 20 min at room temperature. After washing with PBS for 5 min twice, TRAP was stained with NaK tartrate and naphthol AS-BI phosphoric acid. Photomicrographs were taken with a stereomicroscope with a digital camera (Discovery V12 and AxioCam; Carl Zeiss, Inc.). The number of osteoclasts with more than three nuclei was counted and analyzed by GraphPad Prism 6 in a double-blinded manner.

### Resorption pit staining

Mature osteoclasts grown on cortical bovine bone slices were fixed with 4% paraformaldehyde/PBS for 20 minutes. After washing in PBS for 5 min twice, cells were removed from bone slices with a soft brush. The slices were then incubated with 20µg/ml peroxidase-conjugated WGA lectin for 60 min at room temperature. After washing in PBS twice, bone chips were incubated with 0.52 mg/ml 3,3_-diaminobenzidine and 0.03% H2O2 for 30 min. Samples were mounted with 80% glycerol/PBS and photographed with Zeiss AxioPlan2 microscope equipped with an Olympus DP73 digital camera. The resorbed area/bone slice was quantified by ImageJ software (National Institutes of Health), and the percentage of pit area *versus* that of the whole bone slice was calculated and analyzed by GraphPad Prism 6.

### Fluorescent staining of actin filaments and nuclei

Osteoclasts cultured on glass coverslips or bone slices were fixed with 4% paraformaldehyde in PBS for 20min and permeabilized with 0.1% Triton X-100/PBS for 10min at room temperature. The filament actin and nuclei were labeled by Alexa-488 conjugated phalloidin (1:100 from a 1mg/ml stock) and Hoechst 33342 (1:4000 from a 10mg/ml stock), respectively, for 15min at room temperature. After two times 5min-wash with PBS, samples were mounted with 80% glycerol/PBS and photographed under a Zeiss AxioImager Z1 fluorescent microscope equipped with Zeiss AxioCam MRm monochromatic and MR5c color cameras with a set of fluorescent filters. The percentage of active osteoclasts (podosome-belt bearing osteoclasts on glass coverslips and actin-ring bearing osteoclasts on bone slices) out of total osteoclasts was calculated and analyzed by GraphPad Prism 6.

### RNA isolation and real-time qPCR

Total RNA was purified using RNeasy mini kit (Qiagen) according to the manufacture’s protocol. First-strand cDNAs were synthesized from 0.5-1μg of total RNA using the High Capacity cDNA Reverse Transcription kits (Thermo-Fisher Scientific) following the manufacturer’s instructions. TaqMan quantitative real-time PCR was performed using the following primers from Thermo-Fisher Scientific: *Acp5* (Mm00475698_m1); *Nfatc1* (Mm00479445_m1); Mfsd7c (Flvcr2, Mm01302920_m1); *Mrps2* (Mm00475529_m1); *Ppargc1b* (Mm00504720_m1); *Slc11a2* (Mm00435363_m1); *Slc39a14* (Mm01317439); *Slc46a1* (Mm00546630_m1); *Tfrc* (Mm00441941_m1); *Tfr2* (Mm00443703_m1). Samples were amplified using the StepOnePlus real-time PCR system (Life Technologies) with an initial denaturation at 95 °C for 10 min, followed by 40 cycles of 95 °C for 15 s and 60 °C for 1 min. The relative cDNA amount was calculated by normalizing to that of the mitochondrial gene Mrps2, which is steadily expressed in both BMMs and osteoclasts, using the ΔCt method.

### Immunoblotting

Cells were washed with ice-cold PBS twice and lysed in 1 x RIPA buffer containing cOmplete Mini EDTA-free protease inhibitor cocktail. After incubation on ice for 30 min, cell lysates were clarified by centrifugation at 14,000 rpm for 15 min at 4°C. 10 to 30μg of total protein were subjected to 8% or 10% SDS-PAGE gels and transferred electrophoretically onto PVDF membrane by a semi-dry blotting system (Bio-Rad). The membrane was blocked in 5% fat-free milk/Tris-buffered saline for 1 hour and incubated with primary antibodies at 4°C overnight followed by horseradish peroxidase conjugated secondary antibodies. After rinsing 3 times with Tris-buffered saline containing 0.1% Tween 20, the membrane was incubated with ECL for 5 min.

### 59Fe-transferrin uptake and colorimetric total cellular iron measurement

Mouse apo- transferrin was labeled with ^59^Fe and gel-filtered on a Sephadex G-50 column. For Tf-dependent ^59^Fe influx measurements, 25µg of ^59^Fe-labeled Tf was added to each well of a 6-well plate with 3 ml of medium and incubated in a CO2 incubator (37°C, 5% CO_2_) with rotation at 80 rpm. At various times after adding labeled Tf, the medium was aspirated, and cells were washed gently three times with 2 ml of cold PBS. Wells were extracted with 1 ml of 0.1 N NaOH; radioactivity was determined on a 0.5-ml aliquot by a gamma counter. Protein (Bio-Rad) was determined on 25µl of the extract.

To measure cellular iron, cells cultured in 6-well plates were collected in 50µl of the Iron Assay Buffer provided in an Iron Assay kit (catalog no. ab83366, Abcam) and were homogenized with pellet pestles associated with a motor. The cells lysates were centrifuged at 15,000 rpm for 10 min.

The supernatants proceeded to measure the total iron (Fe^2+^ and Fe^3+^) concentration following the manufacturer’s protocol.

### Quantitative proteomic and mass-spectrometry

Cells were harvest and lysed in 2% SDS lysis buffer. A total of 100µg of proteins in each cell lysate were reduced, alkylated, and digested using filter-aided sample preparation (FASP). Tryptic peptides were cleaned by solid phase extraction (SPE), normalized and labeled using a TMTsixplex^TM^ isobaric Mass Tagging kit (catalog no. 90064, Thermo-Fisher Scientific). The labeled peptides were cleaned by SPE and mixed. The mixed peptides from each cell culture were separated into 36 fractions on a 100 x 1.0 mm Acquity BEH C18 column (Waters) using an UltiMate 3000 UHPLC system (Thermo-Fisher Scientific) with a 40 min gradient from 99:1 to 60:40 buffer A:B (Buffer A contains 0.5% acetonitrile and 10 mM ammonium hydroxide. Buffer B contains 10 mM ammonium hydroxide in acetonitrile) ratio under basic (pH 10) conditions and then consolidated into 13 super-fractions.

Each super-fraction was then further separated by reverse phase XSelect CSH C18 2.5 mm resin (Waters) on an in-line 150 x 0.075 mm column using an UltiMate 3000 RSLCnano system (Thermo). Peptides were eluted using a 60 min gradient from 97:3 to 60:40 buffer A : B ratio. Here, buffer A contains 0.1% formic acid and 0.5% acetonitrile and buffer B contains 0.1% formic acid and 99.9% acetonitrile. Eluted peptides were ionized by electrospray (2.15 kV) followed by mass spectrometric analysis on an Orbitrap Fusion Lumos mass spectrometer (Thermo) using multi-notch MS3 parameters. MS data were acquired using the FTMS analyzer in top-speed profile mode at a resolution of 120 000 over a range of 375 to 1500 m/z. Following CID activation with normalized collision energy of 35.0, MS/MS data were acquired using the ion trap analyzer in centroid mode and normal mass range. Using synchronous precursor selection, up to 10 MS/MS precursors were selected for HCD activation with normalized collision energy of 65.0, followed by acquisition of MS3 reporter ion data using the FTMS analyzer in profile mode at a resolution of 50 000 over a range of 100–500 m/z.

Proteins were identified and reporter ions quantified by searching the UniprotKB Mouse database using MaxQuant (version 1.6.10.43, Max Planck Institute) with a parent ion tolerance of 3 ppm, a fragment ion tolerance of 0.5 Da, a reporter ion tolerance of 0.001 Da, trypsin/P enzyme with 2 missed cleavages, variable modifications including oxidation on M and acetyl on protein N-term, and fixed modification of carbamidomethyl on C. Protein identifications were accepted if they could be established with less than 1.0% false discovery. Proteins identified only by modified peptides were removed. Protein probabilities were assigned by the Protein Prophet algorithm. TMT MS3 reporter ion intensity values were analyzed for changes in total protein.

### Mitochondrial mass, ROS production, and membrane potential measurements

Mitochondrial content, mitochondria-derived ROS, and mitochondrial membrane potential were measured using MitoTracker Green fluorescence (catalog no. M7514), MitoSOX^TM^ Red Mitochondrial Superoxide Indicator (catalog no. M36008), and JC-1 Dye (catalog no. T3168) from Thermo-Fisher Scientific, respectively, using fluorescence microscopy. Briefly, cells cultured onto glass-bottom imaging dishes (catalog no. P35G-1.5-14-C, MatTek) were incubated with MitoTracker Green (50nM), MitoSOX (5µM), or JC-1 (5µg/ml), respectively, for 15 min at 37 °C. The probes were then washed off, and cells were examined under fluorescent microscope. Images were analyzed using ImageJ software.

### Seahorse mitochondrial flux analysis

Bone marrow monocytes were plated in wells of Seahorse XF96 cell culture plates. The cells were cultured with 10ng/ml M-CSF alone or 10ng/ml M-CSF plus 100 ng/ml recombinant RANKL for 2 days to generate monocytes and pre-osteoclasts or 4 days for mature osteoclasts. On the day of respiratory function measurements, the media in the wells were changed to unbuffered Dulbecco’s modified Eagle’s medium supplemented with 4 mM glutamate and incubated in a non-CO_2_ incubator for 1 hour at 37°C. Three baseline measurements were acquired before injection of mitochondrial inhibitors or uncouplers. OCR measurements were taken after sequential addition of oligomycin (10µM), FCCP (5µM), and rotenone/antimycin A mixture (10µM). Oxygen consumption rates were calculated by the Seahorse XF-96 software and represent an average of three measurements on 18–24 different wells. The rate of measured oxygen consumption was reported as fmol of O_2_ consumed per min per cell.

### Statistics

Based on power analysis using standard deviation (SD) and the population variance estimated from our previous studies and reports by others, the required sample size of mice for α = 0.05 (two-sided) and a power of 0.95 is ≥ 6 animals per group. The *in vivo* biological replicates are defined as an individual mouse for each experiment. All *in vitro* data were representatives of individual biologic replicates from independent experiments and not technical replicates (repeated measurements of the same sample). For all graphs, data are represented as the mean ± SD. For comparison of 2 groups, data were analyzed using a 2-tailed Student’s *t* test. For comparison of more than 2 groups, data were analyzed using 1-way ANOVA, and the Bonferroni procedure was used for Tukey comparison. For all statistical tests, the analysis was performed using Prism 6 (GraphPad Software, La Jolla, CA) and a *P* value of less than 0.05 was considered significant.

### Study approval

All animal protocols and procedures used in animal studies were approved by the Institutional Animal Care and Use Committees of the University of Arkansas for Medical Sciences, Long Beach VA Healthcare System, and Loma Linda VA Healthcare System. The protocols for generation and use of recombinant DNAs and retroviruses were approved by Institutional Biosafety Committee of the University of Arkansas for Medical Sciences.

## Author contributions

BKD, LW, TF, JZ, designed and performed experiments in conditional knockout mice and *in vitro* osteoclast assays. They participated in analyzing data and preparing the manuscript. NAB and KJK designed and performed mitochondrial experiments. NAB interpreted and analyzed data and edited the manuscript. RL, SGM, RE performed quantitative proteomic experiment and analyzed mass spectrometry data. MLJ designed and performed transferrin uptake and iron assays and helped in editing the manuscript. XW and JQF made substantial contributions to bone histology. TB largely contributed to conditional knockout mice and edited the manuscript, JG, AK, LG performed gene expression and biochemical experiments. LG helped in experimental design and manuscript preparation. WX performed, analyzed, and interpreted the µCT and histological studies. SM and HZ financially supported and conceived the project, supervised experimental design and data analysis, wrote the manuscript with help of all coauthors.

## Acknowledgements

We would like to thank Dr. Nancy C Andrews and Dr. Takashi Nakamura for kindly providing *Tfr1*-flox mice and *Ctsk*-Cre mice, respectively. Erin Hogan at the University of Arkansas for Medical Sciences and Neil T Hoa at the Long Beach VA Healthcare System are acknowledged for help with microscopes. We thank Nancy Lowen and Sheila Pourteymoor for technical assistance. NAB and KJK were supported by NIH grant P20 GM 109005. The University of Arkansas for Medical Sciences Proteomics Core is supported by NIH grants R24GM137786 and P20 GM121293. LG was supported by the Research Scholar Grant funded by the American Cancer Society and the VA Merit Award. SM is a recipient of a Senior Research Career Scientist Award from the Department of Veterans Affairs and has grant support from NIH/NIAMS R01AR048139 and AR070806. The work was supported by grants from NIH/NIAMS R01AR062012, R21AR068509, R01AR073298.

---

**Supplemental Figure 1.**
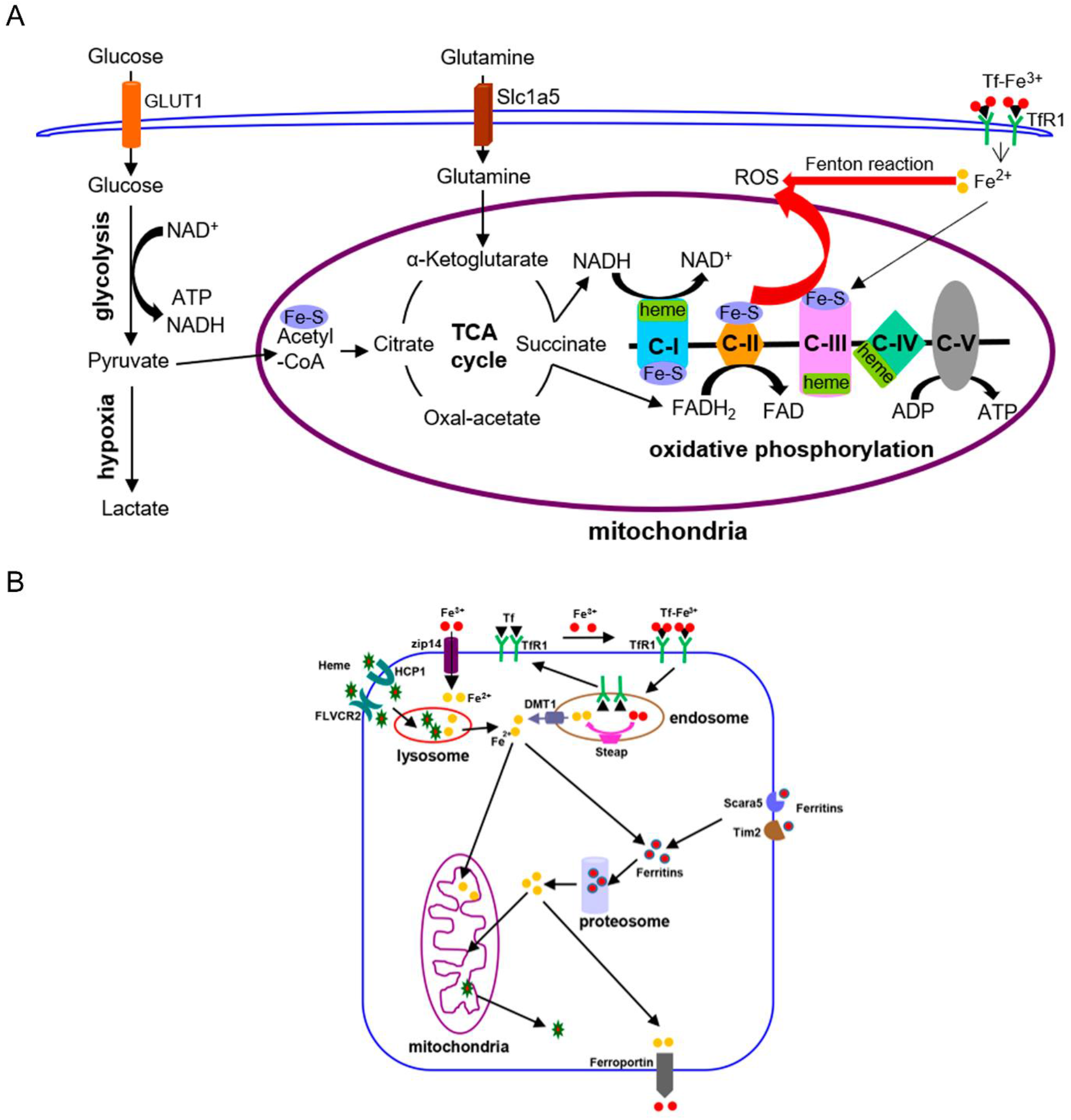
Graphic diagrams of iron in energy metabolism in mammalian cells (A) and cellular iron-uptake pathways (B). C-I to C-V, mitochondrial respiration chain complex-I to complex-V; DMT1, divalent metal ion transporter 1; GLUT1, glucose transporter 1; ROS, reactive oxygen species; Slc1a5, glutamine transporter; Steap, metalloreductase of the six transmembrane epithelial antigen of the prostate family proteins; Tf, transferrin; TfR1, transferrin receptor 1.

**Supplemental Figure 2.**
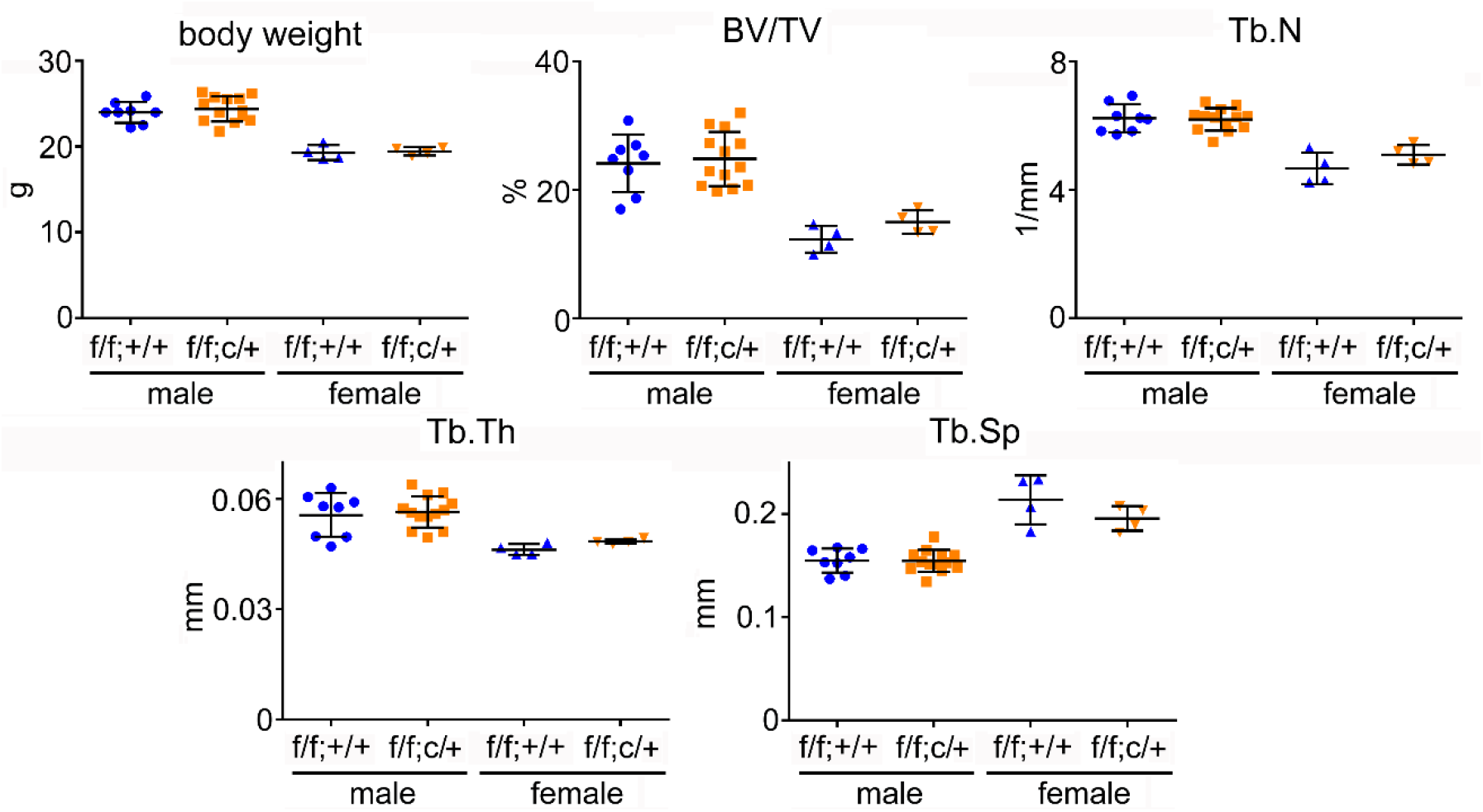
Partial deletion of Tfr1 by one-allele of LysM-Cre has no effects on bone mass and structure in C57BL6/J male and female mice. µCT analysis of trabecular bone mass and structure of distal femurs from C57BL6/J background mice. Tb, trabecular bone; BV/TV, bone volume/tissue volume; Tb.N, trabecular number; Tb. Th, trabecular thickness; Tb.Sp, trabecular spacing. n = 4-13.

**Supplemental Figure 3.**
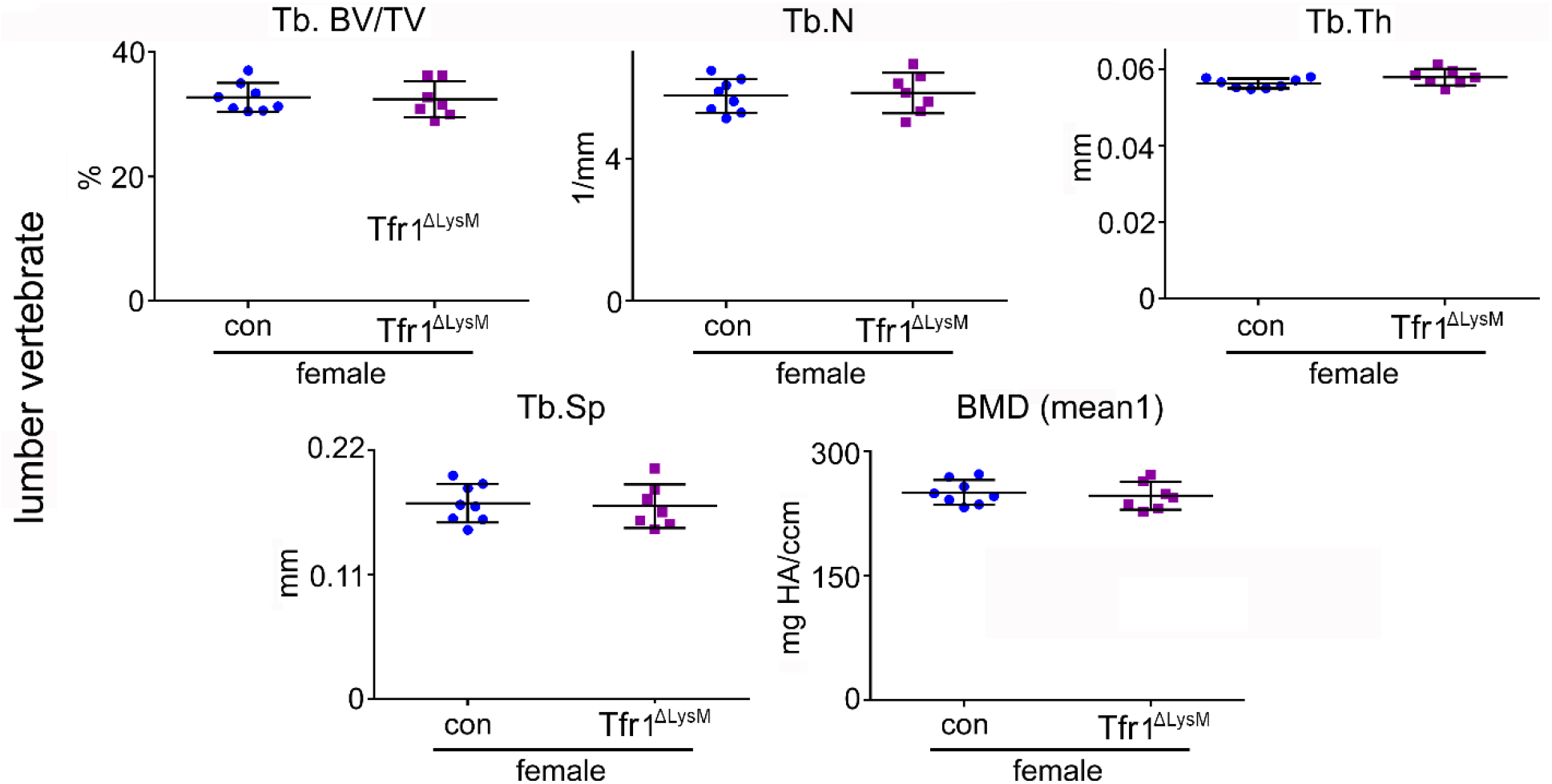
Loss of Tfr1 in myeloid osteoclast precursor cells had little effects on trabecular bone mass and density of lumber vertebrae of 10-week old female mice. µCT analysis of lumber vertebrae. Tb, trabecular bone; BV/TV, bone volume/tissue volume; Tb.N, trabecular number; Tb. Th, trabecular thickness; Tb.Sp, trabecular spacing; BMD, bone mineral density; Cort, cortical bone. * p < 0.05 vs con by one-way ANOVA. n = 7-8.

**Supplemental Figure 4.**
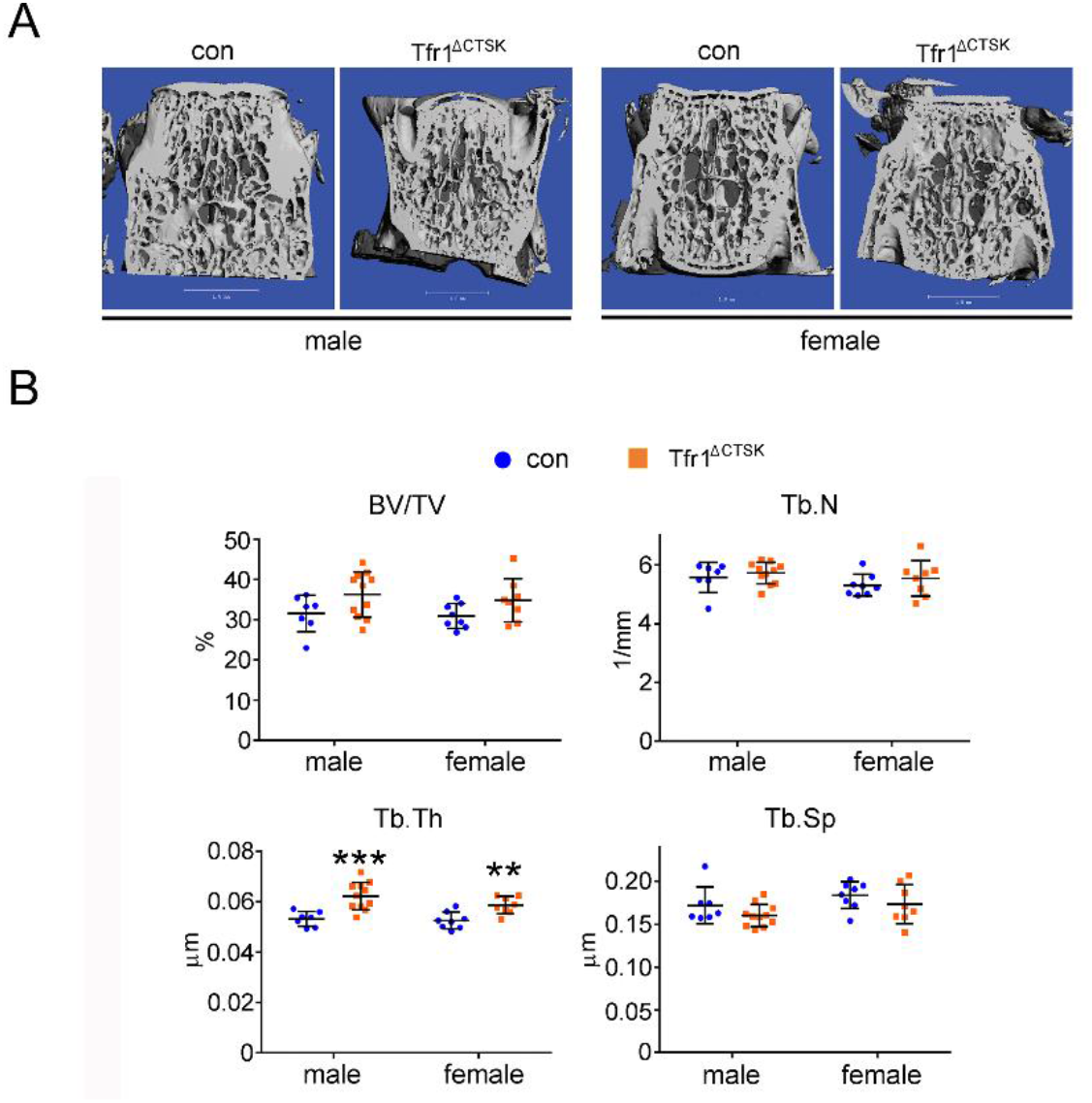
Deletion of Tfr1 in mature osteoclasts slightly increases trabecular thickness of lumber vertebrae of 10-week old mice in mixed background. (**A**) Representative µCT images of L4 lumber vertebra of male and female control (con) and conditional knockout (Tfr1^ΔCTSK^) mice. (**B**) µCT analysis of bone mass and structure of L4 lumber vertebra. Tb, trabecular bone; BV/TV, bone volume/tissue volume; Tb.N, trabecular number; Tb. Th, trabecular thickness; Tb.Sp, trabecular spacing. ** p < 0.01, *** p < 0.001 vs con by one-way ANOVA. n = 7-11.

